# Molecular Basis of Ionic Suppression of ZAP-70 Dependent T Cell Receptor Activation

**DOI:** 10.1101/2025.04.25.650559

**Authors:** Swarnendu Roy, Soumee SenGupta, Kaustav Gangopadhyay, Jibitesh Das, Anushka Sinha, Sudipta Majumder, Prosad Kumar Das, Manas Pratim Chakraborty, Samiran Mondal, Bidisha Sinha, Amirul Islam Mallick, Rahul Das

**Author notes:** Corresponding authors Rahul Das, Amirul Islam Mallick. Author contributed equally (Swarnendu Roy, Soumee SenGupta).

## Abstract

Ionic imbalance in the tumor microenvironment alters the tumor-infiltrating T lymphocyte function. High extracellular K^+^ suppresses T cell function by negatively regulating T cell receptor (TCR) signaling. In contrast, elevated extracellular Na^+^ enhances T cell effector function by boosting the phosphorylation of TCR signaling modules. Here, we presented a mechanism explaining how the two monovalent cations differently regulate TCR function. At rest, high intracellular K^+^ uncouples allosteric recruitment of ZAP-70, a key signaling module, to the TCR complex. The formation of antigen TCR complex induces K^+^ efflux, causing spontaneous recruitment of ZAP-70 to the TCR. Increasing extracellular K^+^ perturbs K^+^ efflux and slows ZAP-70 recruitment to the TCR complex, even upon antigen binding. This leads to defects in T cell development and arthritis-like symptoms in juvenile mice. We conclude that K^+^ dynamics is integral to T cell ligand discrimination and fundamental to turning off the signaling during T cell quiescence.

## Introduction

Monovalent salts of sodium (Na^+^) and potassium (K^+^) ions are the key regulators of T cell effector function in the tumor microenvironment (TME) ^1–5^. Counter-intuitively, K^+^ regulates T cell effector function in the TME differently than Na^+^. The elevated Na^+^ enhances the effector function of tumor-infiltrating T lymphocytes (TIL), whereas high K^+^ in the TME suppresses the TIL function. Both the monovalent cations influence TIL function by altering cell metabolism and T cell receptor (TCR) signaling. For example, elevated K^+^ induces functional caloric restriction of TIL by limiting glucose uptake while promoting T cell stemness ^1^. Additionally, high extracellular K^+^ in TME suppresses TCR signaling by dephosphorylating downstream signaling modules ^1,2,6^. In contrast, elevated Na^+^ aids phosphorylation of TCR signaling modules and glutamine consumption, boosting the effector function of CD8^+^ T cells ^3,4^. Moreover, a high-salt diet (containing NaCl) promotes CD4^+^ T cell differentiation to T helper 17 (Th17). It impairs T regulatory (T_reg_) cell’s suppressive function, linking high dietary sodium to various autoimmune disorders ^7,8^. The molecular basis of why Na^+^ and K^+^ antithetically regulate TCR signaling is an open question.

Engagement of TCR with the antigen presented through the major histocompatibility complex (MHC) of the antigen-presenting cells initiates T cell signaling (Figure 1A) ^9,10^. Antigen binding induces restructuring of the TCR microcluster, forming an immune synapse (IS) ^11,12^. Initially, two tyrosine kinases, a Src family kinase Lck and a Syk family kinase ZAP-70, are recruited to the TCR ^13^. Lck phosphorylates multiple tyrosine residues in the immunoreceptor tyrosine-based activation motifs (ITAM) in the CD3 chains ^13,14^. The ZAP-70 is then spontaneously recruited to the TCR microcluster by binding to the doubly phosphorylated tyrosine residues in the ITAM (ITAM-Y_2_P) motifs ^15–17^. Subsequent autophosphorylation of a tyrosine residue in the activation loop of the kinase domain activates the ZAP-70 ^18,19^. The activated ZAP-70 then phosphorylates scaffold protein LAT that, in turn, recruits PLCγ, eventually turning on the downstream Ca^2+^ influx, Akt-mTOR, and Erk signaling pathways (Figure 1A) ^9,13^. Two potassium channels, shaker-related voltage-gated potassium channels (Kv1.3) and Ca^2+^ activated potassium channel (K_Ca_3.1), are the primary regulator of membrane potential and calcium flux in T lymphocytes ^20–22^. The Kv1.3 colocalized to the TCR microcluster within minutes of antigen binding ^20,23–25^. The K^+^ efflux thus helps sustain Ca^2+^ influx and propagation of TCR signaling downstream ^24,26–28^. Selective blocking of Kv1.3 channels suppresses the immune response and inhibits anti-CD3-dependent T cell proliferation, supporting a fundamental role of intracellular K^+^ in T cell physiology ^21,29,30^. The potassium channel blockers are thus effective therapeutic molecules to treat diverse autoimmune diseases like rheumatoid arthritis (RA), multiple sclerosis, and Type-1 diabetes ^31–34^. However, the molecular mechanism of how intercellular K^+^ regulates TCR activation and ligand discrimination remains elusive.

**Figure 1:**
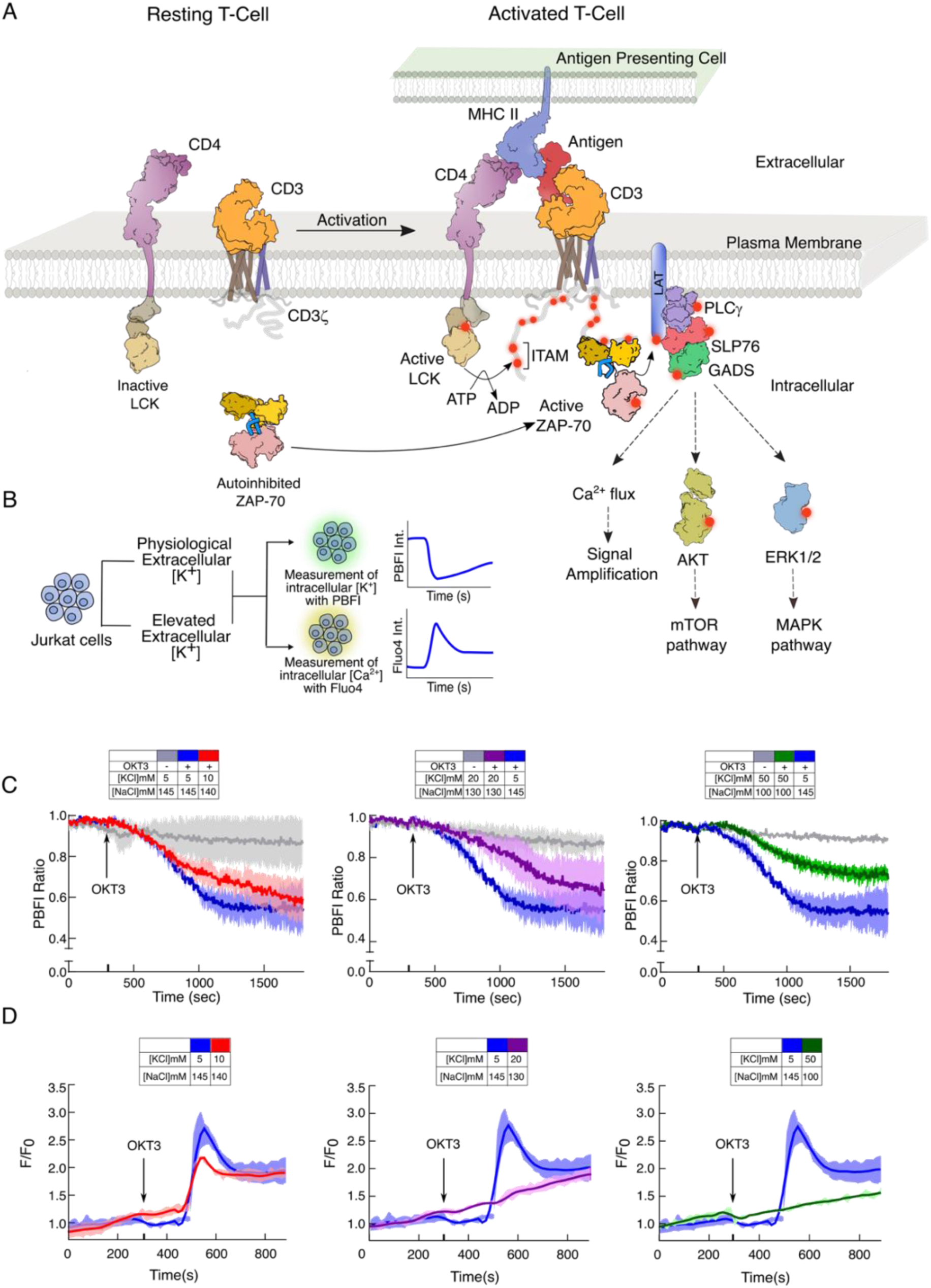
Measurement of TCR-induced potassium efflux and calcium influx in activated Jurkat E6.1 T cells. A) A schematic representation of the TCR signaling pathway. The signaling modules in the TCR complex are labeled. B) The experimental outline for estimating intracellular potassium and calcium levels in Jurkat E6.1 T cells. The T cells are activated with an anti-CD3 antibody (OKT3) in different extracellular KCl concentrations. The intracellular potassium level is measured from the ratio of PBFI fluorescence intensity measured at λ_ex_ of 340 nm and 380 nm, respectively, and λ_em_ of 505 nm. The intracellular calcium level is determined from the change in fluorescence intensity of Fluo4 dyes when measured at λ_ex_ of 495 nm and λ_em_ of 516 nm. C) The potassium efflux is measured in the Jurkat E6.1 T cells at the indicated extracellular KCl concentration. The T cells were activated with an anti-CD3 antibody (OKT3) at the indicated time. The change in PBFI fluorescence intensity is plotted as a function of time. D) The cytosolic calcium level is determined from the plot of normalized Fluo4 intensity as a function of time. At the indicated time, the T cells were activated in the presence of indicated KCl concentration with OKT3. The Fluo4 intensity for the activated (F) T cells is normalized against the Fluo4 intensity measured for the inactive (F_o_) cells. In panel C-D, the solid lines represent the mean from three independent experiments, and the area fill denotes the SD. All data were plotted using GraphPad PrismVer9.5.1. The schematics and icons were made using Inkscape Ver 1.4. See supplementary Figure S1.

Kinetic proofreading assists T cells in discriminating between self and non-self-antigen by calibrating TCR response to the antigen binding ^35^. The half-life of the antigen-receptor complex, the subsequent delay in LAT phosphorylation by ZAP-70, and the time taken for switching Ca^2+^ influx are crucial for TCR selectivity and sensitivity ^36–39^. The short-lived self-antigen-TCR complex will be disassembled before the downstream enzymes are recruited to the IS. At the same time, only the long-lived antigen-TCR complex will initiate a downstream TCR response. How the ionic imbalance interferes with the kinetic proofreading of TCR is unknown.

This paper studied the TCR response after antigen binding in varying salt compositions. We presented a molecular mechanism explaining how ionic imbalance uncouples antigen binding from downstream signaling. We observed that the K^+^ efflux in response to TCR stimulation is sensitive to extracellular salt composition. Our *in vivo* (animal) experiments reveal that elevated serum potassium manifests RA-like symptoms in juvenile mice, resembling the phenotype in ZAP-70^W165C^ mutated SKG mice ^40^. An elevated extracellular K^+^ slows ZAP-70 recruitment to the IS, switching off the Ca^2+^ influx and downstream signaling. Our data suggest that K^+^ is a key allosteric regulator that, at higher intracellular concentrations, uncouples ZAP-70 ligand binding at the TCR complex. We proposed that the intracellular K^+^ dynamics are integral to TCR ligand discrimination and maintain the basal signaling off.

## Results

### High extracellular potassium prevents K^+^ efflux and Ca^2+^ influx upon TCR stimulation

At rest, the negative membrane potential of the T cell is sustained by maintaining a high intracellular [K^+^] concentration (∼ 130 mM) ^1,41^. Engagement of TCR and antigen induces Ca^2+^ influx, resulting in cell membrane depolarization. The negative membrane potential is re-established by effluxing K^+^ out of the cell, mainly by Kv1.3 channels ^20,23^. An increase of extracellular [K^+^] to 20-40 mM in the TME (from ∼ 5 mM [K^+^] in serum) marginally increases intracellular [K^+^] concentration (140 −150 mM) in TIL ^1^. How ionic imbalance in the microenvironment modulates K^+^ efflux and Ca^2+^ influx upon T cell stimulation is lacking, mainly due to the absence of direct measurement of K^+^ efflux.

Here, we measure K^+^ efflux and Ca^2+^ influx in activated Jurkat E6.1 T cells in media containing various ionic compositions (Figure 1B-D). The change in intracellular [K^+^] or [Ca^2+^] concentrations was determined from the change in PBFI-AM ^42,43^ or Fluo4 ^44^ fluorescence intensity, respectively, after activating the T cells with anti-CD3-monoclonal antibody (OKT3). A decrease in PBFI-AM fluorescence intensity indicates K^+^ efflux, whereas an increase in Fluo4 fluorescence intensity indicates Ca^2+^ influx. We observed that stimulation of Jurkat E6.1 cells decreases intracellular [K^+^] concentration and induces Ca^2+^ flux (Figures 1C-D). Under physiological [K^+^] concentration (5 mM), K^+^ efflux begins immediately after TCR activation, reaching a steady state in about eleven minutes (Figure 1C). We noticed a delayed start for the Ca^2+^ influx, and the signal spiked within five minutes of TCR activation (Figure 1D). However, higher extracellular [K^+^] concentration perturbs the TCR-dependent K^+^ efflux and attenuates subsequent Ca^2+^ signaling. At a KCl concentration of 20mM or higher, the Ca^2+^ flux is completely attenuated (Figures 1D and S1C). Higher extracellular K^+^ reduces the rate of K^+^ efflux and maintains a relatively higher intercellular K^+^ level compared to the T cells activated under physiological K^+^ levels (Figures 1C and S1D-E). We ask if maintaining a higher intracellular potassium concentration can decouple the antigen binding from downstream Ca^2+^ signaling.

Kv1.3 potassium channel is essential for antigen-dependent T lymphocyte activation and proliferation ^20,23,30^. Thus, blocking the Kv1.3 channels with clofazimine ^45^ will prevent K^+^ efflux, retaining high intracellular potassium levels even after TCR activation (Figure S1). Inhibiting the Kv1.3 channel prevents Ca^2+^ influx upon antigen binding (Figure S1A) ^22,26^, indicating that high intracellular K^+^ may suppress T lymphocyte function by decoupling the antigen binding and Ca^2+^ flux. It is no wonder why several Kv1.3 inhibitors that suppress T cell function have the potential to treat multiple autoimmune disorders ^31–34,46,47^. However, prolonged treatment with Kv1.3 blockers has previously been shown to inhibit the thymic development of T lymphocytes ^29^. We speculate that intracellular K^+^ dynamics may be fundamental to regulating T cell function and development. We wonder if higher serum K^+^ levels would trigger T cell dysfunction.

### Elevated serum potassium causes RA-like symptoms in juvenile mice

To investigate the role of potassium in maintaining overall immune health, we generated hyperkalemia in three-week-old weaned male BALB/c mice. We selectively blocked the Renin-Angiotensin-Aldosterone System (RAAS) in mice by oral gavaging with a combination of Enalapril (an ACE inhibitor), Amiloride (a potassium-sparing diuretic), and 1.2% KCl solution ^48,49^. The mice were treated with the drug combination for twenty-eight days, as summarised in Figure 2A. We recorded the physiological and behavioral changes throughout the treatment period. At the end of the treatment period, mice were euthanized, and samples were collected for serum potassium estimation, histopathology of joints, and immunophenotyping.

**Figure 2:**
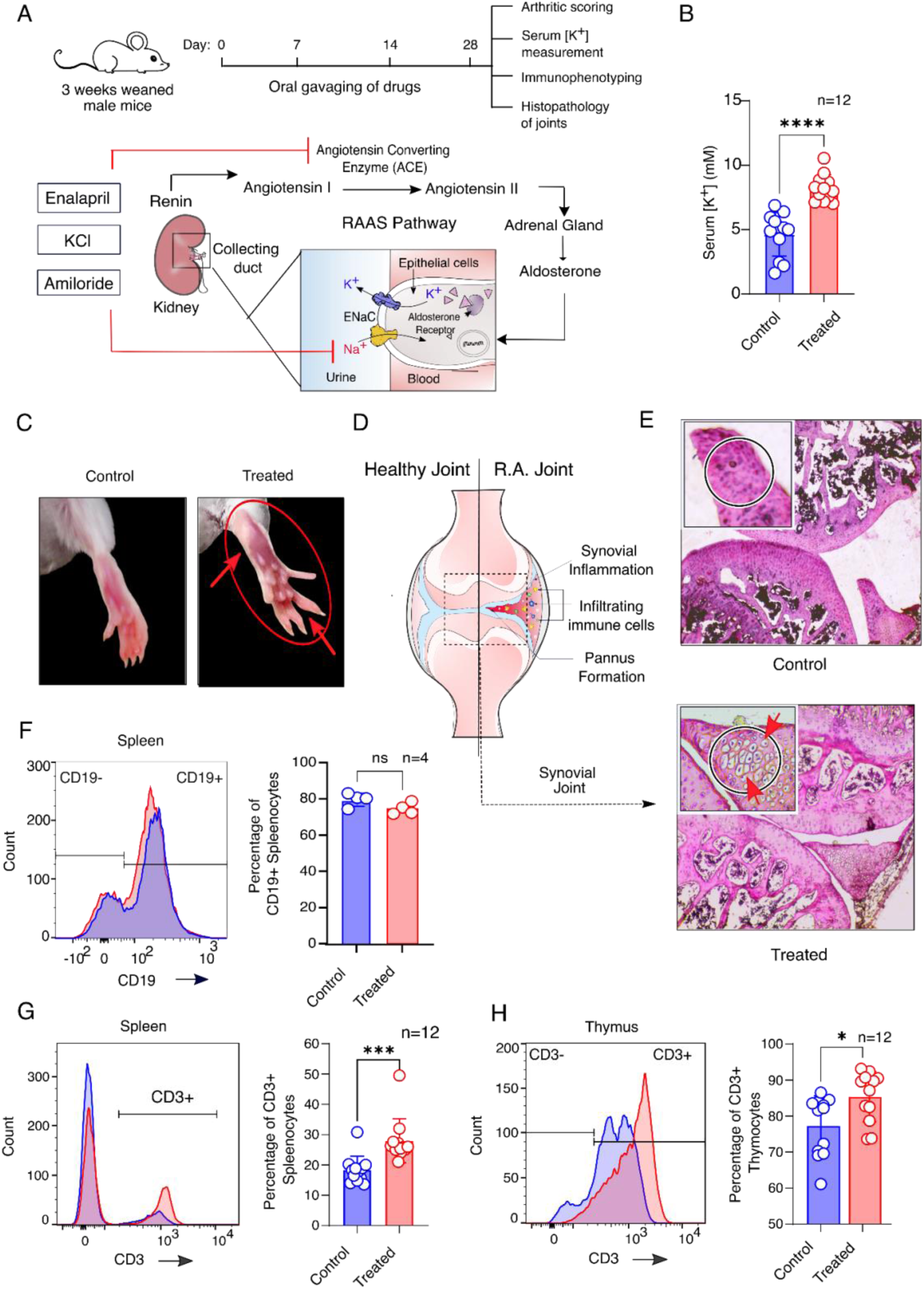
Elevated serum potassium causes RA-like symptoms in juvenile mice. A) Schematic representation of the *in-vivo* animal study for inducing hyperkalemia in juvenile BALB/c mice. Three weeks old weaned male BALB/c mice were orally gavaged with a combination of Enalapril, Amiloride, and 1.2% KCl solution, consecutively for 28 days. The mice were sacrificed and evaluated for serum potassium levels and autoimmunity. B) Comparative analysis of serum potassium concentration in the control (blue) and treated (red) group of mice after 28 days of drug treatment (n=12), *P*<0.0001. C) Representative images of hindlimbs from the experimental mice (control and treated). The swollen joints in the hindlimb of the treated mice are marked with red circles and arrows. D) Cartoon representation of a healthy synovial joint (on the left) and a Rheumatoid arthritic (RA) joint (on the right). The characteristic pannus formation, synovial inflammation, and infiltration of immune cells in the RA joint are labeled. E) Haematoxylin and Eosin (H&E) staining of the synovial tissue of the knee joint from the experimental mice (at 10X magnification). The inset represents the zoomed (40X) cross-section for the control (top) and treated (bottom). In the treated mice, the pannus formation and the infiltrating inflammatory cells are indicated with red arrows. F) Histogram (on the left) showing the total B cell count in splenocytes stained with anti-CD19(PE) mAb. The bar graph (on the right) shows the percentage of total CD19^+^ B cells in the spleen of control (blue) and treated (red) groups (n=4), *P*=0.0848. G) and H) Histogram showing the total T cell count in splenocytes and thymocytes stained with anti-CD3(FITC) mAb, respectively. The adjacent bar graphs in each panel show the percentage of total CD3^+^ T cells in the spleen (G) and thymus (H) of control (blue) and treated (red) mice, respectively, (n=12), G) *P*=0.0008; H) *P*=0.0169. B; F-H) A statistical analysis of two-tailed Students’ t-tests was performed. The data represents mean ± SD (ns= not significant; \**P*<0.05; \*\**P*<0.01; \*\*\**P*<0.001; \*\*\*\**P*<0.0001). All data were plotted using GraphPad PrismVer9.5.1. The flow cytometry data were analyzed using FlowJo Ver8. The schematics and icons were made using Inkscape Ver1.4. See Supplementary Figure S2.

Administrating mice orally with the drug combination and KCl markedly elevated the serum [K^+^] concentration to 8.2 ± 0.29 mM, compared to 4.6 ± 0.47 mM in the control group. The treated mice demonstrated a phenotype resembling autoimmune Rheumatoid Arthritis (RA) reported in the SKG mice ^40^. For example, we observed progressive development of inflammatory lesions in the joints of the hindlimbs, forelimbs, and digits (fingers), with significantly higher arthritic scores in the treatment group (Figures 2C and S2D). Additionally, loss of body hair, hunchback, and reduced grooming behavior, indicate possible alterations in the immune system of the treated group of mice (Figures S2A-B) ^50^. The radiographic and histopathological examination of the treated mice revealed narrowed joint spaces with thickened synovial tissue, pannus formation, and infiltration of proinflammatory cells suggestive of advanced joint inflammation resembling arthritis-like conditions (Figure S2C and 2D-E). As anticipated, the treated group of mice exhibited impaired limb movement and related pain sensitivity in the affected limbs. In addition to autoimmune arthritis-related pain, we assume that the pain sensation may be associated with impaired neurotransmission resulting from elevated serum K⁺ levels.

T cells in the SKG mice cannot discriminate between self and non-self-antigens due to a mutation in ZAP-70 that prevents kinase activation upon antigen binding ^35,40^. The ZAP-70^W165C^ mutation alters the thymic differentiation of CD3^+^ T cells by tweaking the kinetic proofreading ability of the TCR complex, resulting in a positive selection of autoreactive T cells ^40,51,52^. Since hyperkalemia in juvenile mice induces autoimmune RA-like symptoms, we ask if increasing serum potassium level also affects T cell development and function.

### Elevated serum potassium perturbs T cell development and function

We evaluate the total T and B lymphocyte count in the control and hyperkalemic mice. We observed no difference in the CD19^+^ B cell in the splenocytes between the control and treated group, suggesting elevated serum potassium does not alter the total B cell population (Figure 2F). In contrast, we observed an increase in CD3^+^ T cell count in the thymocytes and splenocytes of hyperkalemic mice compared to the control group (Figures 2G-H). We wonder if the imbalance in the T cell population and RA-like symptoms in hyperkalemic mice is because of defects in T cell development.

Next, we probed the effect of elevated serum K^+^ on the thymic maturation of CD3^+^ T cells. We evaluated the critical checkpoints in T cell development by immune profiling the Double Negative (DN), CD4^+^CD8^+^ Double Positive (DP), or CD4^+^/CD8^+^ Single Positive (SP) stages (Figures 3 and S3) ^53^. We used a combination of CD44^+^ and CD25^+^ markers to differentiate between DN stages (Figure 3A) ^54^. In the thymus of the hyperkalaemic mice, we observed an overall increase in the DN cell count and a reduction in the number of DP cells (Figure 3B). However, we did not observe a significant change in the number of CD4^+^ or CD8^+^ SP thymocytes and splenocytes (Figure 3B and S3E) between the treatment and control groups of mice. Differential staining of various DN stages reveals a decrease in DN1 thymocyte count. At the same time, we observe a significant increase in the DN3 and DN4 cell populations (Figure 3A and S3B).

**Figure 3:**
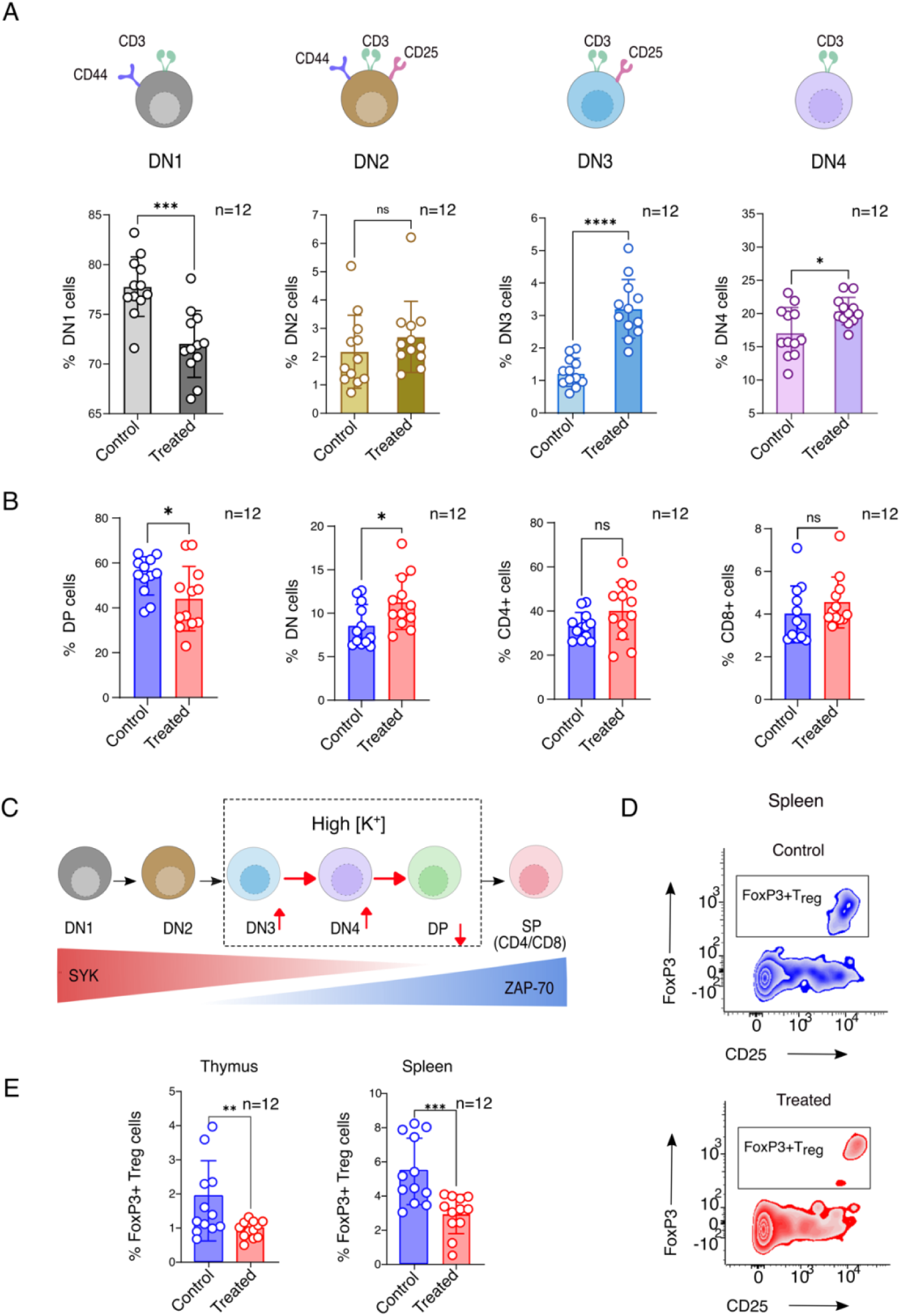
Elevated serum potassium perturbs T cell development. A) Bar graphs compare the percentage (%) of cells in each Double Negative (DN) stage of developing T thymocytes in the control and treated groups of mice. The top panel depicts the cartoon representation of the indicated DN stage markers used in the flow cytometric separation of the thymocytes. (n=12), DN1 *P*=0.0002; DN2 *P*=0.3245; DN3 *P*<0.0001; DN4 *P*=0.0167. B) Bar graphs comparing the percentage of thymocytes in the Double Negative (DN), Double Positive (DP), CD4^+,^ and CD8^+^ Single Positive cells in the control and treated groups of mice (n=12), DN *P*=0.0273; DP *P*=0.0464; CD4^+^ *P*=0.1063; CD8^+^ *P*=0.2922. C) Schematic representation summarizing the effect of elevated potassium on T cell development. The expression patterns of Syk family kinases during the T cell development are marked below ^58^. D) Representative flow cytometric analysis to determine the frequencies of CD4^+^regulatory T cells (T_reg_) in the spleen. The T_reg_ cells are labeled with anti-CD3, anti-CD4, anti-CD25, and anti-FoxP3 mAb. E) Bar graph comparing the percentage of T_reg_ cells in the thymus (on the left) and spleen (on the right). The control and treated groups of mice are colored blue and red, respectively. (n=12), T_reg_(thymus) *P*=0.0088 and T_reg_(spleen) *P*=0.0001. A; B; E) A statistical analysis of two-tailed Students’ t-tests was performed. The data represents mean ± SD (ns= not significant; \**P*<0.05; \*\**P*<0.01; \*\*\**P*<0.001; \*\*\*\**P*<0.0001). All data were plotted using GraphPad PrismVer9.5.1. The flow cytometry data were analyzed using FlowJo Ver8. The schematics and icons were made using Inkscape Ver 1.4. See Supplementary Figure S3.

Defects in the number or function of regulatory T (T_reg_) cells are primarily linked to various autoimmune disorders like type 1 diabetes, multiple sclerosis, systemic lupus erythematosus, and rheumatoid arthritis ^55^. Recent studies connect high-salt-diet-induced T_reg_ cell dysfunction to autoimmunity, suggesting a pivotal role of salt balance in regulating T_reg_ cell function. Our flow cytometric analysis showed a significant reduction in CD4^+^CD25^+^ FoxP3^+^ T_reg_ cells in both the thymus and spleen of hyperkalemic mice (Figures 3D, E, and S3F). Since T_reg_ cells play a crucial role in suppressing the immune response against self-antigens ^56,57^, we conclude a decrease in T_reg_ cell count due to high serum potassium levels may have resulted in autoimmune RA-like symptoms in the treated mice.

To summarize, increased serum potassium profoundly affects the DN3 to DP stages of thymocyte maturation (Figure 3C). The expression of two Syk family tyrosine kinases, Syk and ZAP-70, are dynamically regulated during the thymic development of T cells ^53^. Syk and ZAP-70 expression overlaps in the DN3 and DN4 stages. Impaired activation of either kinase may arrest the T cell maturation ^58^. Moreover, ZAP-70 is indispensable for sustained pre-TCR/TCR signaling from DN4 to DP stages ^59^ and also for positive and negative selection of DP cells to self-tolerant single positive CD4^+^ or CD8^+^ naïve T cells ^60^ (Figure 3C). Thus, in our experiments, impediment of thymocyte maturation in the DN3 to DP stages and reduction in the T_reg_ cell population indicates that elevated serum potassium may impair ZAP-70 function.

### High extracellular potassium interferes with the ZAP-70-dependent TCR signaling

TCR does not possess intrinsic catalytic activity. The signaling starts with the sequential recruitment of two tyrosine kinases, Lck and ZAP-70 (Figure 4A) ^13^. Thus, we begin by probing the role of excess extracellular K^+^ on overall phosphorylation status in activated Jurkat E6.1 T cells. We observed higher extracellular potassium-impaired total tyrosine phosphorylation levels of Jurkat E6.1 T cells compared to those activated under physiological [K^+^] concentration (Figure S4A). Our data indicates that potassium negatively regulates tyrosine kinase activation in a concentration-dependent manner. Therefore, we next focus on the effect of elevated K^+^ on the phosphorylation of key tyrosine residues in the activation loop of Lck (Y394) and ZAP-70 (Y493). The phosphorylation of tyrosine in the activation loop is a hallmark of the activated kinase ^9^. The phosphotyrosine stabilizes the active conformation of the enzyme by preventing the activation loop from folding back into the catalytic site ^61^. We observed that elevated extracellular K^+^ inhibits ZAP-70 activation in a concentration-dependent manner (Figures 4B-C). The phosphorylation of Y493 was significantly reduced at 20 mM KCl compared to the physiological concentration of 5 mM. However, elevated extracellular potassium did not inhibit Lck Y394 phosphorylation (Figures 4B-C), suggesting ZAP-70 is specifically inhibited at higher [K^+^]. As anticipated, inhibiting ZAP-70 at higher [K^+^] reduces the phosphorylation of multiple downstream signaling modules, like LAT, PLCγ, Erk, and Akt (Figures 4B-D and S4B). To independently verify if higher intracellular K^+^ inhibits ZAP-70 activation, we determined the Y493 phosphorylation in activated Jurkat E6.1 T cells treated with clofazimine (Figure S4C). Clofazimine maintains higher intracellular [K^+^] by blocking K^+^ efflux through Kv1.3 channels during TCR activation (Figures S 1B, D, and E). Clofazimine treatment significantly reduced ZAP-70 autophosphorylation even after TCR stimulation, suggesting high intracellular K^+^ inhibits ZAP-70 activation.

**Figure 4:**
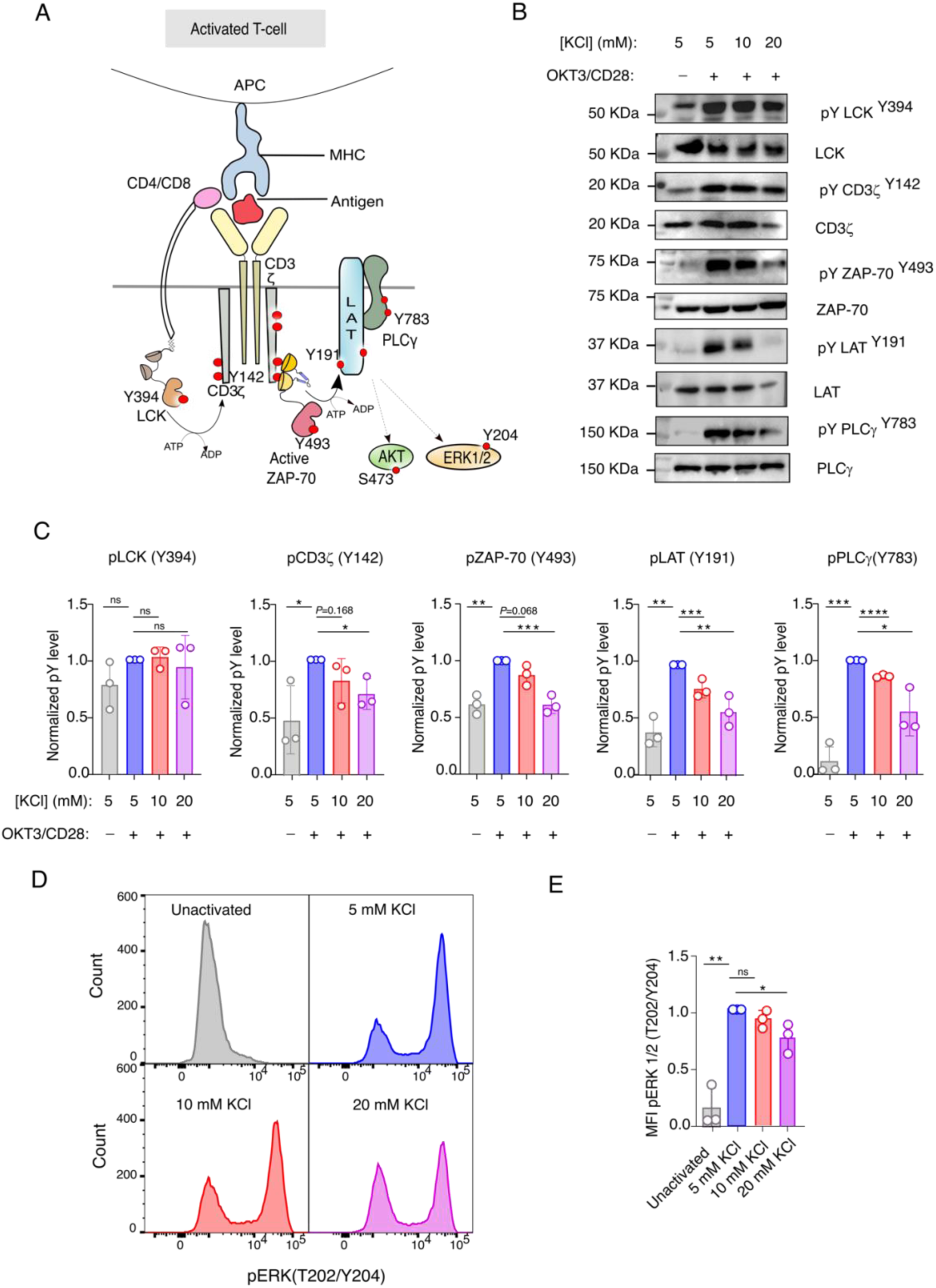
Intracellular potassium levels regulate ZAP-70-dependent TCR signaling. A) Schematic representation of the activated TCR signaling pathway. The signaling modules evaluated in this study are labeled. B) Representative immunoblot showing the phosphorylation of various TCR signaling modules. The Jurkat E6.1 T cells were stimulated with OKT3/anti-CD28 antibodies in the indicated concentration of extracellular KCl and immunoblotted with indicated anti-pY and anti-protein antibodies. C) Densitometric analysis of the immunoblots shown in panel B. Bar graphs represents the fold changes in phospho-tyrosine levels of specific proteins in the TCR signaling modules at the indicated potassium concentration. For pLCK (Y394) from left to right: *P*=0.1306; *P*=0.6623; *P*=0.7016, for pCD3ζ (Y142) from left to right: *P=*0.0373; *P=*0.1680; *P=*0.0175, for pZAP-70 (Y493) from left to right: *P*=0.0014; *P*=0.0688; *P*=0.0010, for pLAT (Y191) from left to right: *P*=0.0017; *P*=0.0002; *P*=0.0012, for pPLCγ (Y783) from left to right: *P*=0.0003; *P*<0.0001; *P*=0.0219. D) Representative flow cytometry histograms of phospho-ERK1/2 (T202/Y204) in the unactivated and activated Jurkat E6.1 T cells at the indicated extracellular potassium concentration. E) Bar graphs showing the fold changes of ERK 1/2 phosphorylation in stimulated Jurkat cells at the indicated potassium concentration in the extracellular media. For pERK (T202/Y204) from left to right: *P*=0.0012; *P*=0.1268; *P*=0.0312. C; E) A statistical analysis of two-tailed Students’ t-tests was performed. Each data represents the mean ± SD from three independent experiments. (ns= not significant; \**P*<0.05; \*\**P*<0.01; \*\*\**P*<0.001; \*\*\*\**P*<0.0001). All data were plotted using GraphPad PrismVer9.5.1. The flow cytometry data were analyzed using FlowJo Ver8. The schematics and icons were made using Inkscape Ver 1.4. See Supplementary Figure S4.

Since higher intracellular potassium does not perturb Lck activation, it is surprising that elevated extracellular K^+^ reduces the phosphorylation of tyrosine residues in the ITAM motifs of the CD3-ζ chain (Figure 4B-C). Previous independent studies showed impaired interaction between ZAP-70^W165C^ and CD3-ζ chain in the SKG mice, resulting in reduced CD3-ζ phosphorylation ^40,62^. We speculate that the decreased tyrosine phosphorylation of CD3-ζ is due to the dephosphorylation of ITAM motifs by phosphatases present in the IS, which is otherwise preserved by the ZAP-70 and ITAM-Y_2_P complex ^63^. Therefore, we next investigate how higher intracellular K^+^ concentration prevents ZAP-70 and CD3-ζ interaction.

### Potassium allosterically regulates the interaction between ITAM-Y_2_P in the CD3 chain and the regulatory module of ZAP-70

ZAP-70 bridges a key kinetic bottleneck between the antigen-TCR complex and the LAT-based signalosome essential for ligand discrimination ^37^. The recruitment of ZAP-70 to the plasma membrane is mediated by the interaction between ITAM-Y_2_P in the CD3-ζ and the regulatory module of ZAP-70 (Figure 5A) ^17,63^. The ZAP-70 regulatory module is made of a tandem Src homology domain (tSH2) connected to the C-terminal kinase domain (Figures 5A and B). In the autoinhibited state, the tSH2 domain adopts an open conformation, rendering the regulatory module incompetent to bind the ITAM-Y_2_P (Figure 5B) ^64^. The binding of ITAM-Y_2_P remodels the tSH2 domain structure to a closed conformation, subsequently releasing the autoinhibition of the kinase domain ^53,63^. We first investigate how increased intracellular K^+^ levels modulate the ZAP-70 and CD3 interaction at the plasma membrane. We measured the localization of the ZAP-70 tagged to EGFP to the plasma membrane using total internal reflection fluorescence (TIRF) microscopy following T cell stimulation (Figures 5C-D and S5). The TIRF image shows the formation of distinct EGFP clusters at the plasma membrane of the live Jurkat P116 cells activated in 5 mM and 20 mM KCl (Figures 5C and S5E). Our image analysis revealed that the ZAP-70 clustering is significantly slower when T cells are activated in media containing 20 mM KCl compared to 5 mM KCl (Figure 5D). We ask how higher intracellular potassium perturbs ZAP-70 recruitment to the TCR.

**Figure 5:**
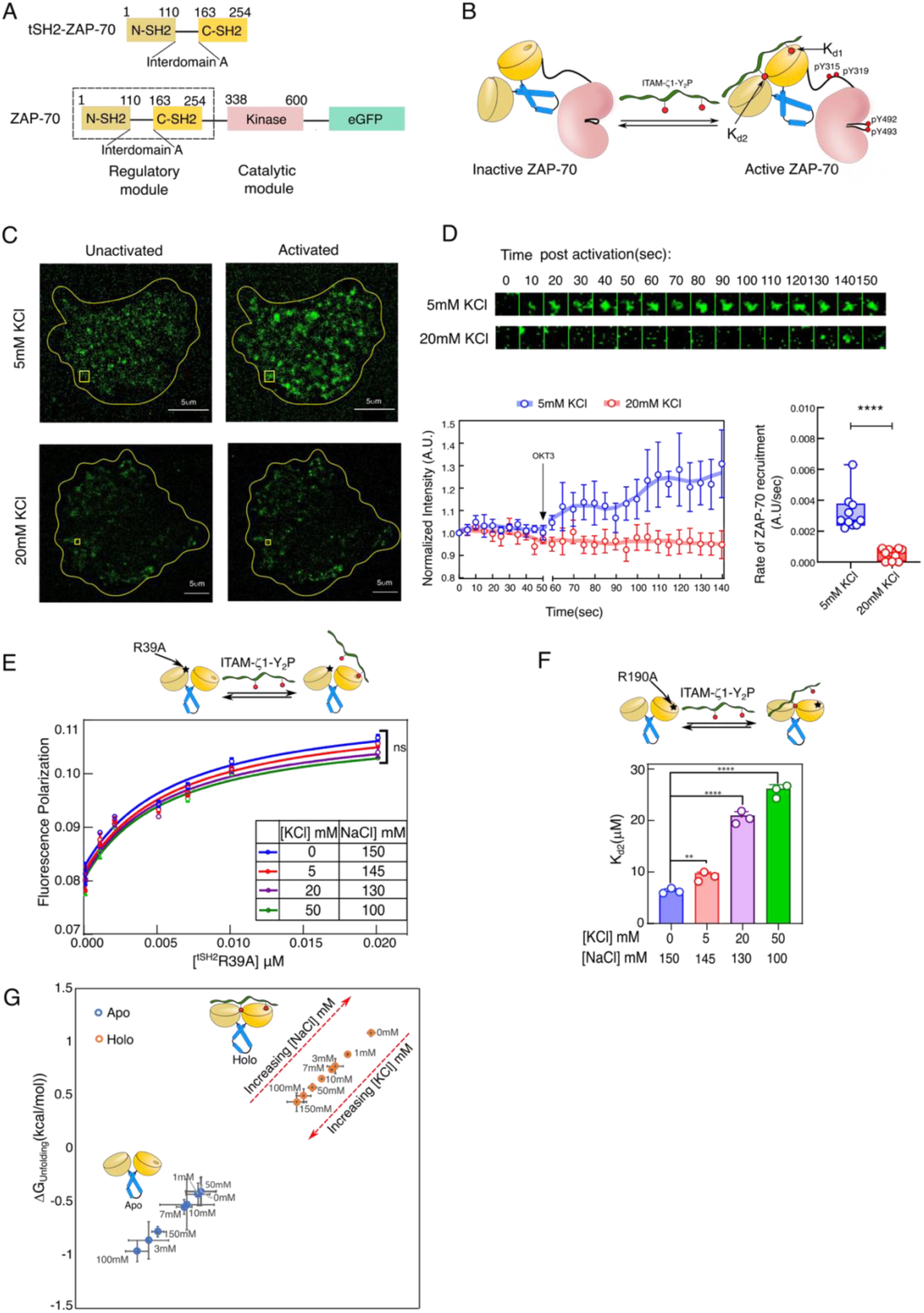
Higher potassium concentration prevents ZAP-70 and CD3-ζ interaction in the TCR complex. A) A schematic representation of the domain architecture of ZAP-70 constructs was used in this study. B) Cartoon representation of ZAP-70 activation model. C) Representative TIRF microscopic image of unactivated (on the left) and activated (on the right) live Jurkat P116 T cell stably expressing ZAP-70-EGFP construct. The cells are activated with anti-CD3/anti-CD28 mAb at the indicated KCl concentration. The cell boundary is marked with a yellow line, and a yellow box indicates a region of the ZAP-70-EGFP cluster. D) The top panel represents a time-lapse montage of an individual ZAP-70-EGFP cluster after activation with OKT3/anti-CD28 mAb at the indicated extracellular KCl concentration. The bottom panel at the left is the plot of EGFP intensity as a function of time, measured from five ZAP-70 clusters (size: 10 pixels, 1pixel= 0.65μm) in one live Jurkat P116 T cell. The arrow indicates the time point when OKT3/ anti-CD28 mAb is added. Each point represents the average intensity of five ZAP-70 clusters, and the error bar denotes the S.D. The red and blue solid lines are guiding lines. The bottom panel on the right is the plot of the rate of change in EGFP intensity from the individual cells at the indicated KCl concentration. The error represents SD from ten cells (n=10), *P*<0.0001. E) The plot of fluorescence polarization as a function of tSH2 domain concentration, determined from the titration of the ^R39A^tSH2 mutant and ITAM-ζ1-Y_2_P tagged to Alexa Fluor488. Each data point is the mean ± SD from three independent experiments. The colored solid lines represent the fitting to the first-order binding. F) The bar graph shows the *K_d_* for the N-SH2 phosphate binding pocket and ITAM-ζ1-Y_2_P interaction. The *K_d_* represents the mean value from three independent titrations of the indicated ZAP-70 construct and ITAM-ζ1-Y_2_P using ITC. *P* value: 5mM KCl: =0.0023, 20mM KCl:<0.0001, 50mM KCl:<0.0001 G) ΔG_unfolding_ for the *apo* and *holo* tSH2 domain of ZAP-70 is plotted at the indicated KCl concentration. The ΔG_unfolding_ is derived from the thermal denaturation profile measured using Circular Dichroism (CD) spectroscopy. Each data point represents the mean ± SD from three independent experiments. D) and F) a statistical analysis of two-tailed Students’ t-tests was performed. Central values and error bars represents mean ± SD (ns= not significant; \**P*<0.05; \*\**P*<0.01; \*\*\**P*<0.001; \*\*\*\**P*<0.0001). All data were plotted using GraphPad PrismVer9.5.1. The image analysis was done using Fiji Ver 1.54m. The schematics and icons were made using Inkscape Ver 1.4. See Supplementary Figures S5 and S6.

The tSH2 domain of ZAP-70 is made of two SH2 domains, N-SH2 and C-SH2 domains, connected by a helical linker (Figure 5A). We have previously shown that the ITAM-Y_2_P binds first to the C-SH2 phosphate binding pocket (PBP) with a strong (nM) affinity (*K_d1_*), forming an encounter complex ^39,62^. That, in turn, induces the tSH2 domain to adopt a closed conformation, allowing phosphotyrosine to bind at the N-SH2 PBP with weaker (μM) affinity (*K_d2_*) (Figure 5B) ^62^. To determine how elevated K^+^ alters ZAP-70 recruitment to TCR, we re-evaluate the steady-state binding of ITAM-Y_2_P to C-SH2 (*K_d1_*) or N-SH2 (*K_d2_*) PBP with increasing KCl concentration (Figure 5E-F). We studied the binding of ITAM-ζ1-Y_2_P and C-SH2 PBP using a tSH2^R39A^ mutant by measuring the fluorescence polarization of an ITAM-ζ1-Y_2_P peptide labeled with AlexaFluor488 (Figures 5E-S6A). We measured the ITAM-ζ1-Y_2_P and N-SH2 PBP interaction by isothermal calorimetric titration (ITC) using a tSH2^R190A^ mutant (Figure 5F and S6D). Our data suggests that increasing potassium does not perturb the binding of ITAM-ζ1-Y_2_P and C-SH2 PBC (Figure 5E). Unexpectedly, potassium perturbs the phosphotyrosine binding to N-SH2 PBP in a concentration-dependent manner (Figure 5F). We observed that the N-SH2 PBP binds to the ITAM-ζ1-Y_2_P with a *K_d2_* = 6.33±0.34 µM in the presence of 150 mM NaCl (Figure 5F and Table S5). In contrast, increasing KCl concentration to 20 mM reduces the binding affinity of N-SH2 PBP for the ITAM-ζ1-Y_2_P by more than three times (*K_d2_* = 20.93±0.82 µM). We noticed that 5 mM KCl marginally affects the N-SH2 PBP and ITAM-ζ1-Y_2_P interaction (*K_d2_* = 9.77±0.61µM) compared to the NaCl (Figure 5F). Our biochemical measurements concur with cell-based studies explaining how elevated K^+^ prevents ZAP-70 clustering at the membrane (Figure S5 C-E). Further analysis of structural stability (from ΔG_unfolding_) suggests that higher K^+^ destabilizes the ligand-bound *holo*-tSH2 domain in a concentration-dependent manner compared to NaCl (Figure 5G). We wonder why increasing potassium concentration destabilizes the tSH2 *holo*-state.

### Potassium slows down the structural transition of ZAP-70 to an active state in a concentration-dependent manner

In the inactive state, the tSH2 domain of ZAP-70 adopts an open conformation where only the C-SH2 PBP is competent to bind the ITAM-Y_2_P(Figure 6A) ^39,62–64^. The final transition to closed conformation of the *holo*-tSH2 domain requires assembling the aromatic stacking interaction (called thermodynamic brake) that allosterically couples the two SH2 domains (Figure 6A) ^39^. The tSH2 domain binds the ITAM-Y_2_P in multiple binding steps, producing a biphasic ligand binding isotherm (Figure 6B) ^62^. We probed the conformational rearrangement of the tSH2 domain during ITAM-ζ1-Y_2_P titration by measuring the changes in the intrinsic tryptophan fluorescence at steady-state (Figure 6B) ^62^. We observed that increasing [K^+^] concentration by varying the NaCl to KCl ratio remodels the binding isotherm of ITAM-ζ1-Y_2_P and tSH2 domain to a hyperbolic (monophasic) pattern (Figure 6B). At 50mM KCl, the binding isotherm resembles the monophasic ligand binding reported previously for the ^W165C^tSH2 mutant ^62^. The W165 is a key residue in the allosteric network coupling the phosphotyrosine binding to the two PBP. Mutating W165C breaks the allosteric coupling between the C-SH2 and N-SH2 PBP, impairing ZAP-70-mediated TCR signaling (Figure 6A) ^40,62^. We ask why potassium preferentially perturbs the ligand binding to the N-SH2 PBP.

**Figure 6.**
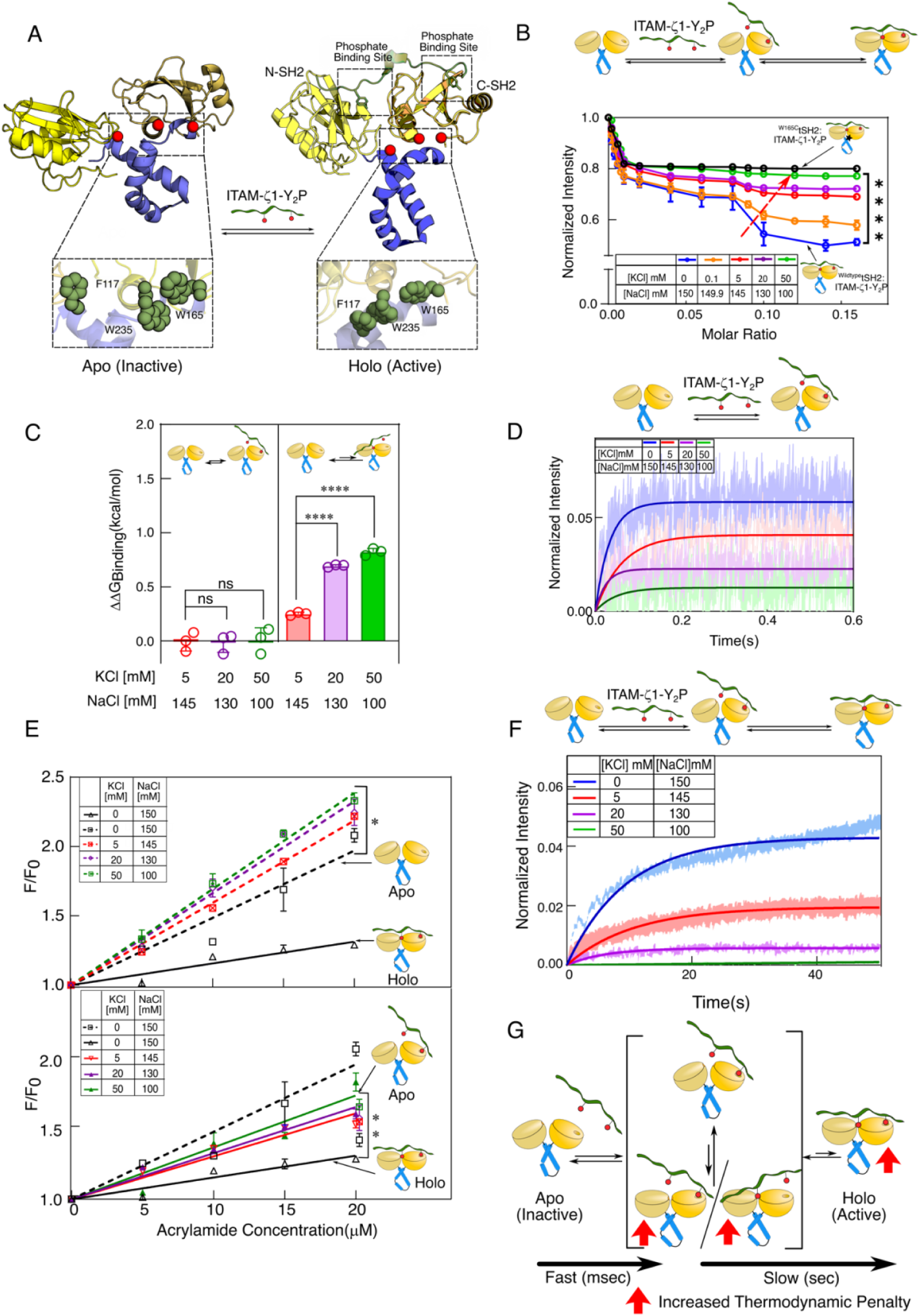
High potassium concentration increases the thermodynamic penalty on the interaction of ITAM-Y_2_P and ZAP-70 tSH2 domains. A) The cartoon structure of the *apo* (PDB ID: 1M61) and *holo* (PDB ID: 2OQ1) ZAP-70 tSH2 domain. Inset highlights the conformation of the aromatic residues in the inactive and active states. B) A plot of changes in ZAP-70 tSH2 domain intrinsic tryptophan fluorescence against ligand to protein molar ratio. ITAM-ζ1-Y_2_P is titrated against the indicated construct of tSH2 domain in the presence of increasing KCl concentration. Each data point represents the mean ± SD from three independent experiments. The solid color lines are guiding lines. The ^W165C^tSH2 mutant found in SKG mice ^40^ is indicated in black. C) The change in ΔG_binding_ (ΔΔG_binding_) for the N-SH2 and C-SH2 phosphate binding pockets is plotted at the indicated KCl concentration. ΔΔG_binding_ is calculated with respect to the ΔG_binding_ obtained in the absence of KCl for the respective phosphate-binding pocket. *P* value from left to right: 0.4442, 0.9051, <0.0001, <0.0001. D) and F) are the plots of time-dependent binding kinetics of the tSH2 domain (100 nM) and ITAM-ζ1-Y_2_P (15 μM) in the fast and slow time scales, respectively. Solid lines represent the single exponential fitting of the blank subtracted data. The area fill represents SD from three independent experiments. E) Stern-Volmer plot of normalized fluorescence intensity against increasing acrylamide concentration for the *apo* (broken lines) and *holo* (solid lines) tSH2 domain at the indicated KCl concentrations. The lines represent the fitting to a straight-line equation. The Stern-Volmer quenching constant (*K_sv_*) was determined from the slope of the fitted lines (see Figure S6). G) A kinetic model of ZAP-70 tSH2 domain and ITAM-Y_2_P interaction ^39^. The red up-arrows indicate the steps where the thermodynamic penalty increases with increasing potassium concentration. C) and E) a statistical analysis of two-tailed Students’ t-tests was performed. Each data represents mean ± SD (ns= not significant; \**P*<0.05; \*\**P*<0.01; \*\*\**P*<0.001; \*\*\*\**P*<0.0001). All data were plotted using GraphPad PrismVer9.5.1. The schematics and icons were made using Inkscape Ver 1.4. See Supplementary Figures S6 and S7.

We recently showed that the initial encounter complex between the ITAM-Y_2_P and C-SH2 domain forms with a fast-kinetic step, which transiently assembles the N-SH2 PBP^39^. The final transition of the tSH2 domain to a closed conformation follows a slow kinetic step. Comparison of change in Gibbs’ free energy for ligand binding 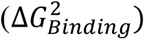 to the N-SH2 PBP shows that higher [K^+^] concentration imparts a greater thermodynamic penalty on ITAM-Y_2_P binding compared to Na^+^ (Figure 6C and Table S5). For instance, compared to the ITAM-ζ1-Y_2_P binding measured at 5 mM KCl (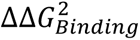 = 0.25±0.012 kcal/mol), 20 mM KCl imposes a significant penalty (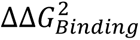 = 0.69±0.009 kcal/mol) on ligand binding to N-SH2 PBP (Figure 6C and Table S5). Suggesting that higher [K^+^] concentrations prevent the assembly of the N-SH2 PBP.

To determine if increasing [K^+^] affects the rate of structural rearrangement upon ligand binding, we measured the kinetics of the encounter complex formation and the final transition closed state. We probed the rate of structural rearrangement to the encounter complex and the final closed state by measuring the binding kinetics of ITAM-ζ1-Y_2_P to the tSH2 domain in the fast 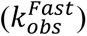 and slow 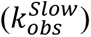 kinetic steps. Using stopped-flow fluorescence spectroscopy, we mixed the tSH2 domain with excess ITAM-ζ1-Y_2_P dissolved in a buffer of various NaCl and KCl concentrations (Figures 6D and F). To probe the fast or slow kinetic step, we recorded the changes in the intrinsic tryptophan fluorescence of tSH2 after ligand mixing for 600 ms or 200 s, respectively. We observed that higher [K^+^] concentration does not alter the fast kinetic step, i.e., the formation of the encounter complex (Figures 6D and S7C). However, the reduced fluorescence intensity observed at higher [K^+^] concentrations indicates destabilization of the transiently formed encounter complex. In contrast, we observed that K^+^ significantly reduced the transition 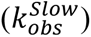 of the tSH2 domain to the final *holo*-state at higher concentrations (Figures 6F and S7D). Suggesting that K^+^ disproportionately affects the structure of the closed conformation of the ligand-bound tSH2 domain.

To probe if elevating K^+^ would perturb the closed conformation of the tSH2 *holo*-state, we determine the Stern-Volmer quenching constant (*K_sv_*) by measuring acrylamide quenching of tryptophan fluorescence at increasing KCl to NaCl ratio. A higher *K_sv_* value for the *apo* tSH2 domain (0.052±0.004 µM^-1^) in buffer containing NaCl suggests that the tSH2 adopts an open conformation (Figure 6E and Table S7). At the same time, shielding of acrylamide quenching of tryptophan fluorescence in the tSH2 *holo*-state (*K_sv_* = 0.0164±0.001 µM^-1^) indicates that the tSH2 domain adopts a closed conformation upon ligand binding, in the presence of NaCl. By increasing KCl concentration, the *K_sv_* increases significantly for the *holo*-tSH2 domain (Figures 6E and S7B). At 50 mM KCl, a *K_sv_* value of 0.042±0.002 µM^-1^ indicates that the tSH2 domain adopts a more open-like conformation even in the ligand-bound state. Our data suggests that higher K^+^ does not perturb the formation of the encounter complex between ITAM-Y_2_P and the C-SH2 domain of ZAP-70 (Figure 6G). However, the higher thermodynamic penalty imposed on the second ligand binding significantly slows the assembly of key aromatic-stacking interaction that allosterically couples structural rearrangement of the tSH2 domain to the active state. We anticipate the impaired aromatic-stacking interaction, in turn, will prevent ZAP-70 activation by slowing its recruitment to the plasma membrane (Figure 6G). Indeed, our studies using Jurkat E6.1 T cells show that increasing extracellular [K^+^] dampens the rate of ZAP-70 activation loop autophosphorylation (Figure 7A). To summarize, our data points towards the central role of the aromatic stacking interaction in the allosteric network in the tSH2 domain for sensing intracellular K^+^ dynamics.

**Figure 7.**
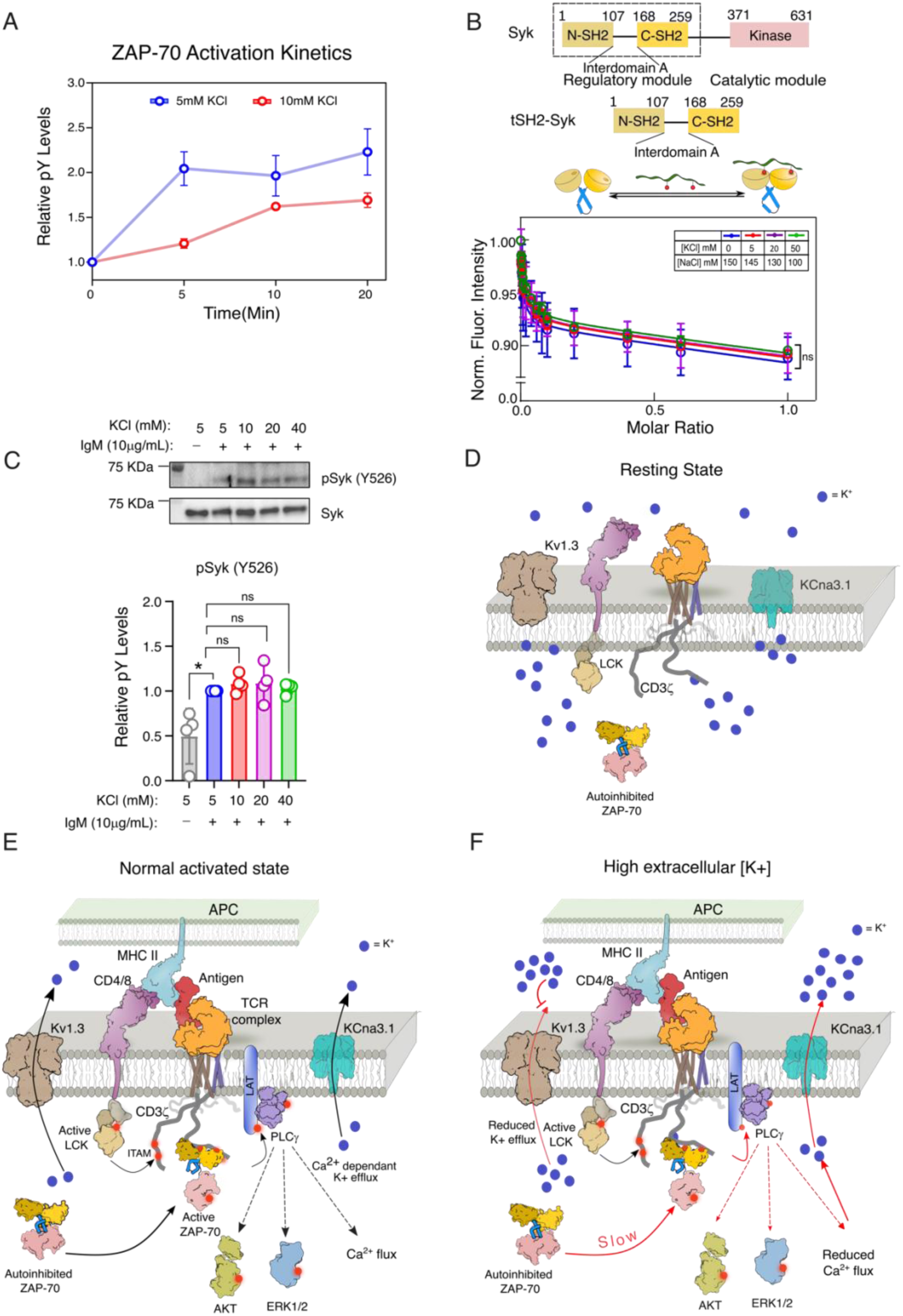
A model of ionic suppression of ZAP-70-dependent early TCR signaling. A) The phosphorylation level of ZAP-70 Y493 upon TCR stimulation is plotted against time. The phosphorylation level is obtained from the densitometric analysis of immunoblot shown in Figure S7E. Each data point represents the mean ± SD from three independent experiments. All data were plotted using GraphPad PrismVer9.5.1. B) The top panel is the schematic representation of the domain architecture of Syk tyrosine kinase used in this study. The bottom panel is the plot of change in intrinsic tryptophan fluorescence in the tSH2 domain of Syk against the ligand-to-protein molar ratio at the indicated salt composition. C) The top panel is the representative immunoblot showing the phosphorylation level of Y526 and expression of Syk in Ramos-RA1 B cell lines. The cells were activated with IgM in the presence of the indicated concentration of extracellular KCl. The bottom panel is the densitometric analysis of the immunoblots shown above. Bar graphs represent the fold changes in phospho-tyrosine levels. From left to right: *P*=0.0170; *P*=0.1529; *P*=0.4141; *P*=0.3127. D) – F) The proposed model of ionic suppression of early TCR signaling. Panel B) represents the resting state of a T cell. Panels C) and D) stimulation of TCR in the absence and presence of elevated extracellular potassium, respectively. The binding of antigen, presented through an APC, to the TCR initiates consecutive recruitment and activation of two tyrosine kinases, Lck and ZAP-70, respectively. Activation of ZAP-70 initiates subsequent downstream signaling. Red arrows indicate the steps perturbed by high extracellular potassium concentration. The schematics and icons were designed using Inkscape Ver1.4. A - C) A statistical analysis of two-tailed Students’ t-tests was performed. Each data represents the mean ± SD from three independent experiments. (ns= not significant; \**P*<0.05; \*\**P*<0.01; \*\*\**P*<0.001; \*\*\*\**P*<0.0001). All data were plotted using GraphPad PrismVer9.5.1. The schematics and icons were made using Inkscape Ver 1.4. See Supplementary Figure S7.

### Syk tSH2 is insensitive to an increase in cellular potassium levels

To further investigate the role of the aromatic stacking interaction in the tSH2 domain of ZAP-70 in sensing the cellular K^+^ levels, we turned to Syk tyrosine kinase expressed in B cells. Syk is the paralogous kinase of ZAP-70 expressed in B cells that couples B cell receptor (BCR) response to the downstream signaling ^65^. ZAP-70 and Syk share a similar structural architecture (Figure 7B) and high sequence homology. Yet, the tSH2 domain of Syk binds to the ITAM-Y_2_P motifs in the BCR in a hyperbolic manner, suggesting the ligand binding is uncooperative ^39,66^. Compared to ZAP-70, ITAM-Y_2_P binding to the Syk tSH2 domain does not rely on the aromatic-stacking interaction (thermodynamic brake) (Figure S7G) ^39^. We speculate that the Syk in B cells will be active even in high extracellular K^+^ levels. Thus, we evaluate the effect of high [K^+^] concentration on the binding of the Syk tSH2 domain to ITAM-Y_2_P and activation of the kinase. We measured the Syk activation loop Y526 phosphorylation in Ramos RA-1 cells after stimulating with IgM in the presence of elevated extracellular [K^+^] concentration. As anticipated, high [K^+^] concentration did not disrupt the tSH2 and ITAM-ζ1-Y_2_P binding (Figures 7B and S7F). We observed no change in Syk activation loop autophosphorylation when Ramos RA-1 cells were activated in increasing [K^+^] concentration (Figure 7C). The differential response of ZAP-70 and Syk to K^+^ suggests that the K^+^ does not influence the ligand binding at the PBP of the tSH2 domains. Instead, K^+^ interferes with the aromatic stacking in the tSH2 domain, thereby uncoupling the structural transition of the encounter complex to the final active state (Figure 6G).

## Discussion

Ion channels and ionic flux mediate diverse physiological functions of lymphocytes in health and disease. In the quiescence state, T cells maintain a high concentration of intracellular K^+^ (Figures 1C and 7B)^1,41^. Early patch-clamp experiments supported the idea that antigen-binding to T cells induces K^+^ efflux, which sustains Ca^2+^ influx (Figures 1C-D and 7B) ^22,24^. We observed that perturbing the gating property of the K^+^ channels by increasing the extracellular [K^+^] concentration ^67^ or blocking potassium channels perturbs TCR-mediated K^+^ efflux and attenuates Ca^2+^ influx (Figures 1C-D and S1A-B) ^22,45^. Here, we presented a molecular mechanism of how intracellular K^+^ levels regulate TCR signaling.

Our data suggest that K^+^ interferes in the early step of TCR signaling. High intracellular K^+^ in the resting state or due to insufficient efflux (in TME) inhibits the recruitment of ZAP-70 to the TCR complex (Figure 7). Intriguingly, gene expression of ZAP-70 and the Kv1.3 channel is modulated by the same epigenetic regulator ^28^. High intracellular K^+^ prevents ITAM-Y_2_P binding to the tSH2 domain of ZAP-70 by imparting a higher thermodynamic penalty on the structural transition to the final closed conformation (Figures 7D and 6G). The structural transition of the tSH2 domain is mediated by a network of allosteric interactions functioning as a thermodynamic brake. Central to the thermodynamic brake are the aromatic-stacking interactions critical for ligand discrimination and sensing cellular K^+^ levels (Figure 6G) ^39^. Our data suggests that K^+^ may prevent the assembly of a key aromatic stacking interaction, possibly by interacting with the π face of the aromatic rings ^68^ (Figure 6A). Thus, increasing [K^+^] concentration alters the ZAP-70 function by imparting a higher thermodynamic penalty on the assembly of the aromatic stacking interaction. That, in turn, puts a brake on the structural transition of the tSH2 domain to the active state (Figure 5B), slowing down the recruitment of ZAP-70 to the IS (Figure 7D). The non-functional thermodynamic brake ^39^ in the tSH2 domain of Syk renders the kinase insensitive to the cellular [K^+^] concentration (Figures 7B-C). Explaining why, in hyperkalemic juvenile mice, increased serum potassium specifically affects T cell count and development in the DN3 to DP stage (Figures 2F-H).

In an aqueous solution, K^+^ is preferred over Na^+^ to form a cation-π interaction with the aromatic residues ^68^. Indeed, in all our biochemical experiments, we observed that Na^+^ facilitates ITAM-Y_2_P interaction with the tSH2 domain. Together, our data explains why Na^+^ and K^+^ differentially regulate TCR signaling ^1,3–5^.

In conclusion, we showed that the intracellular potassium dynamics are critical for regulating the physiological function of the TCR response. High intracellular potassium ensures low receptor activation in the T cell quiescent by preventing ZAP-70 recruitment to the plasma membrane. We proposed that K^+^ allosterically regulates the tSH2 interaction with the ITAM-Y_2_P motif in the CD3 chain at the IS. We conclude that the intracellular potassium dynamics is an integral part of the kinetic proofreading for antigen discrimination by TCR. The concentration dependency of ZAP-70 and ITAM-Y_2_P interaction on intracellular K^+^ levels suggests that the cellular potassium dynamics may fine-tune the TCR response to self and non-self antigens.

## Statistics and reproducibility

Statistical analyses were performed with GraphPad Prism 9 using an unpaired two-tailed student’s t-test. No statistical method was used to predetermine the sample size. No data were excluded from the analyses. All data were presented as mean ± SD from at least three biologically independent experiments. P<0.05 was considered as statistically significant.

## Data Availability Statement

All the relevant data are contained within this article and in the supporting information. Source data are available in the Source Data file and as a Figshare deposition. Uncropped blots are available in the Source Data file.

## Acknowledgments

The authors thank Dr. Tapas Mondal, Professor of pediatrics at McMaster University Canada, for the helpful discussion. The authors thank Tamal Ghosh for helping with the FACS data acquisition and the supporting staff at the Animal House and Analytical Biology facility. The authors thank research funding from IISER Kolkata, infrastructural facilities supported by IISER Kolkata, and DST-FIST (SR/FST/LS-II/2017/93(c)) and DBT builder project (BT/INF/22/SP45383/2022). This work is supported by a grant from DBT (BT/PR54950/BMS/85/671/2024). BS acknowledges support from Wellcome Trust/ DBT India Alliance fellowship (IA/I/13/1/500885) for the TIRF microscope. Fellowships from UGC supported SSG and JD.

## Author Contributions Statement

The manuscript was written through the contribution of all authors. All authors have approved the final version of the manuscript. RD, SR and SSG designed the experiments. RD, AIM, and SSG designed the animal studies. SaM did the blind data analysis of the animal studies. BS, RD, SSG, and JD designed and analyzed the imaging experiments. SR, SSG, KG, MPC, AS, SuM, and PKD performed the Biochemical experiments and data analysis. RD, AIM, SR, and SSG wrote the manuscript.

## Competing Interests Statement

The authors declare that they have no conflict of interest with the contents of this article.

## Notes

### Competing Interest Statement

The authors have declared no competing interest.

