## Supporting information for "Molecular Basis of Ionic Suppression of ZAP-70 Dependent T Cell Receptor Activation"

### Author contributed equally

**Supporting Information**

Corresponding authors

Rahul Das:

Amirul Islam Mallick:

#### Methods

##### Construct and reagents used

The ZAP-70 tSH2 domain (residue number 1-256) cloned into the pSKB2 vector and full-length ZAP-70 was a gift from Prof. John Kuriyan, U.C. Berkeley. The point mutations were introduced into the tSH2 construct (R39A, R190A) using site-directed mutagenesis by PCR, as explained previously [1]. PBFI dye for potassium level estimation and Fluo4 dye for studying calcium flux were procured from Thermo Fischer Scientific (Catalogue: PBFI, AM, Cell Permeant-Special Packaging, #P1265MP) and (Catalogue: Fluo-4 NW Calcium Assay Kit, F36206), respectively. The ITAM-ζ1-$Y_2$P peptide was purchased from S Biochem. The AlexaFluor 488 tagged ITAM-ζ1-$Y_2$P peptide was obtained from GenPro Biotech. Antibodies used in this study are provided in the Table S1.

##### Cell culture

Jurkat E6.1 cells (ATCC # TIB-152), Jurkat P116 cells (ATCC # CRL-2676) and Ramos RA-1(ATCC # CRL-1596) were cultured in high glucose RPMI 1640 media (GIBCO #A10491-01) and HEK293T cells were cultured in DMEM media (GIBCO #11995049), supplemented with 10%FBS (GIBCO #10270-106) and 1% anti-anti (GIBCO#15240-062) at $37^0$C and 5% $CO_2$. Before activation, the cells were washed with PBS. $1 \times 10^6$ cells (or $5 \times 10^6$ as per experiment) were resuspended in serum-free RPMI media for 6 hours at 37°C. Following serum starvation, cells were resuspended in media containing various concentrations of KCl for an additional 4 hours.

##### Intracellular Potassium level estimation in Jurkat T-cells

The serum-starved Jurkat E6.1 T cells (at $5X10^6$ cells/mL ) were loaded with PBFI-AM, probenecid, and Pluronic F127 by incubating for three hrs in RPMI-1640 media at 37 °C and 5% $CO_2$ [2]. Then the cells were washed twice in HBSS (composed of 0.4mM Potassium Phosphate monobasic, 0.3mM Sodium Phosphate dibasic pH 7.4, 0.5mM Magnesium Sulfate Heptahydrate, 0.5mM Magnesium Chloride Hexahydrate, 4mM Sodium Bicarbonate, 6mM Glucose, one mM $MgCl_2$) to remove the excess dye and resuspended in HBSS containing various proportions of KCl and NaCl. The fluorescence intensity of the cells was recorded by excitation at two wavelengths ($\lambda_{ex}$) 340 and 380nm, and the fluorescence intensity was recorded ($\lambda_{em}$) at 505 nm with 5 sec intervals. At the beginning of each experiment, the baseline (blank) was recorded for 5 minutes. The cells were then stimulated with OKT3 (1: 1000 dilution), and the change in PBFI fluorescence intensity was recorded for 30 minutes. In the control (blank) experiment, OKT3 was replaced by an equal amount of HBSS buffer. The intracellular KCl level was determined from the plot of normalized fluorescence intensity measured as a function of time. The PBFI-AM intensity was normalized from the ratio of fluorescence emission recorded at $\lambda_{ex}$ 340 from $\lambda_{ex}$ 380nm.

##### Calcium Signalling in Jurkat T-cells

The serum-starved Jurkat T-cells were treated with Fluo4 and incubated for one hour in HBSS [3-6]. The cells were washed twice in HBSS to remove the excess dye and resuspended in HBSS containing various proportions of KCl and NaCl. The fluorescence intensity of Fluo4 was measured with $\lambda_{ex}$ of 495 nm and $\lambda_{em}$ of 516 nm. At the beginning of each experiment, the baseline was recorded for 300 seconds, and then the cells were activated with OKT3 (1: 1000 dilution). A control (blank) experiment was performed for each condition where an equal volume of HBSS was added instead of OKT3. The calcium flux was estimated from the normalized Fluo4 fluorescence intensity plotted against time. The Fluo4 fluorescence intensity of the activated cells (F) was normalized with respect to the fluorescence intensity of the control (blank) experiment ($F_0$).

**Immunoblot analysis**

The serum-starved Jurkat T-cells were stimulated with 1ug/mL human anti-CD3 mAb (OKT3) and 5ug/mL anti-CD28 mAb for 5mins at 37˚C (or as indicated in the respective figures). Whereas, the Ramos RA-1 cells were activated with 10 µg/ml of IgM for 5 min (REF). The phosphorylation is quenched by adding lysis buffer (final composition of 50 mM Tris pH 8, 150 mM NaCl, 1% NP40, 2 mM $Na_3VO_4$, 1mM Benzamidine,10 mM NaF, 5 mM EDTA, 1x PhosSTOP (Roche #04906845001) [7]. The cell lysate was placed on ice for 30 minutes, and cell debris was cleared by centrifuging at 13,000 × g. Protein samples were prepared by heating the supernatant with NuPAGE LDS sample buffer (4X) (Thermo #NP0007), resolved on SDS-PAGE, and blotted onto a PVDF membrane. The blotted membrane was blocked with 3% BSA dissolved in 1× TBS (20 mM Tris pH 7.6 and 150 NaCl) and 0.1% Tween-20 for 1 hour at RT and then incubated with primary antibody at 4 °C overnight (dilution 1:1000). The unbound primary antibody was washed thrice with 1× TBST containing 0.1% tween-20, followed by incubation with a secondary antibody diluted in 1% BSA (dilution 1: 2500 for both anti-rabbit and anti-mouse HRP secondary antibody). The blot was washed three times with 1× TBST and two times with 1XTBS before developing with the ClarityTM Western ECL substrate kit (Bio-Rad #1705060). All images were acquired using the Bio-Rad Chemidoc system and analyzed using ImageJ. The details of the antibodies used and the dilutions are mentioned in Supplementary Table 1. For potassium channel inhibition, cells were treated with 5µM clofazimine for one hour before stimulation [8].

**Flow Cytometric analysis to study Phospho-protein activation in Jurkat T-cells**

The unstimulated and stimulated (for 5 min) Jurkat cells were immediately fixed in paraformaldehyde (final concentration of 4%) to quench the reaction. The cells were incubated at 25 °C for 30 min, and then washed once with FACS buffer (2% FBS, 1 mM EDTA in PBS buffer), and resuspended in 100% ice-cold methanol for 30 mins at 4°C to permeabilize. The cells were rehydrated with FACS buffer for one hour on ice and stained with specific anti-phosphotyrosine- or anti-protein antibodies for one hour at room temperature. The cells were washed twice in FACS buffer and then stained with a secondary antibody conjugated to Alexa Fluor-488 or AlexaFluor-647 for 45 minutes at room temperature. The cells were washed twice in FACS buffer and analyzed using BD LSRFortessa flow cytometer (BD Biosciences). [7,9].

***In vivo* animal experiments**

For all animal experiments, around three weeks weaned male BALB/c mice were used. The mice were housed in the institute small animal house facility of IISER Kolkata. The animals were maintained and treated as per protocols approved by IAEC, IISER Kolkata, CCSEA, Department of Animal Husbandry and Dairy (DAHD) Govt. of India, (Approval no. IISERK/IAEC/AP/2022/74.01).

##### Hyperkalemia induction

The mice were divided into two groups: the control and the treatment group. The control mice were fed a normal diet. To induce hyperkalemia, mice in the treatment group were orally gavaged with a combination drug (Amiloride Hydrochloride and Enalapril) and KCl solution consecutively for 28 days. The KCl and the drugs were fed as per the following dose: 1.2% KCl solution in water, 0.5mM Amiloride Hydrochloride (a potassium-sparing diuretic) at a dosage of 10ml/Kg/day, and 0.12mg/ml Enalapril (Enam 10) (an ACE inhibitor) at a dosage of 30mg/Kg/day dissolved in drinking water [10-13]. The body weight, drinking water intake, and behavioral changes of the experimental animals were monitored throughout the treatment period.

##### Arthritic Scoring and X-ray of swollen joints

The arthritic score was calculated as described previously [14]. Briefly, the swellings in the limb joints in mice were monitored by observation and scoring as follows: 0 = no swelling; 0.1= swelling of one finger joint; 0.5 = mild swelling of wrist or ankle; 1= severe swelling of wrist or ankle. Scores for all joints were summed for individual mice recorded over an interval for the entire period (twenty-eight days) of drug treatment. The X-ray photographs of the live experimental mice were captured on the day of sacrifice using an *in-vivo* animal imaging system (Spectral Instrument Imaging Model: Ami HTX) at a high-power X-ray mode.

##### Estimation of serum potassium level

At the end of the treatment regime, the animals were euthanized, and the blood was collected by cardiac puncture. The blood was allowed to clot by incubating at room temperature for 15-20 minutes. Straw-colored serum in the supernatant was collected by centrifugation at 1000 x g for 10 minutes at $4^0$C. The potassium concentration in each sample was analyzed using a fluorometric potassium assay kit (ABCAM #ab252904). The serum potassium level was determined following the manufacturer's protocol using GraphPad PrismVer9.5.1.

##### Histopathological studies of joints

To study inflammatory changes in the limbs, the knee joints were dissected for histopathological studies. The tissue samples were fixed in 4% paraformaldehyde for 36–48 hours. The tissues are then processed in increasing concentrations of ethanol and xylene, and embedded in a single mold of paraffin. The embedded tissue samples were then sectioned into 5 µm thick transverse sections and stained with hematoxylin and eosin. The tissues were examined for evidence of inflammations like infiltration of immune cells in the swollen joints, pannus formation, and bone erosion. [15]

##### Immunophenotyping of cell population in Thymus and Spleen

To determine the population of total T-cells, total B-cells, and various subtypes of T-cells ($CD4^+$ SP, $CD8^+$SP, $T_{reg}$, DN, and DP), mononuclear cells were isolated from the thymus and spleens of the experimental mice. Briefly, both groups of mice were sacrificed at the end of the treatment period, and thymus and spleens were collected and washed thrice with PBS. The tissue was homogenized in a sterile petri dish in 1 mL complete RPMI 1640 (GIBCO #A10491-

01) growth media. The cell suspension was filtered using a 70 μm cell strainer, and the mononucleated cells were collected using Histopaque (Sigma-Aldrich, 10771) density gradient. The sample was then processed for flow cytometric analysis [16]. The fluorescently labeled primary antibodies for detecting various mononucleated cells are provided in Table S1.

**Stable cell line preparation by lentiviral transduction**

Human full-length ZAP-70 fused to EGFP at the C-terminal was cloned into second-generation lentivirus vector pLVX M.Puro (Addgene #125839). To make the final lentivirus carrying the ZAP-70-EGFP, the cloned pLVX plasmid is cotransfected along with packaging vector psPAX2 (Addgene #12260) and envelope vector pMD2.G (Addgene #12259) (in 4:3:1 ratio) into HEK293T cells [17]. The supernatants containing the virus were collected 48 hours post-transfection, filtered using 0.45μm filters, and stored at -80$^{O}$C. For stable cell line preparation, Jurkat P116 cells were activated using OKT3/anti- CD28 mAb (1μg/mL and 5μg/mL) for 10 minutes and then transduced with the lentiviral titer and incubated for forty-eight hours at 37 $^{o}$C and 5% $CO_2$. Jurkat P116 cells expressing ZAP-70-EGFP were sorted using BD FACSAria III and cultured in fresh complete RPMI media (GIBCO A1049101) with 15% FBS (GIBCO 10270-106) and 1% anti-anti (GIBCO 15240062).

**Live cell imaging using TIRF microscopy**

Jurkat P116 cells stably expressing ZAP-70-EGFP were incubated for 3 hours in RMPI media containing 5mM KCl or 20mM KCl. For imaging, the cells adhered on a glass coverslip coated with human fibronectin (Sigma-Aldrich F2006) for one hour at 37 $^{o}$C and in 5% $CO_2$. The cells were then imaged using TIRF microscopy with a time-lapse of 5 s/frame, utilizing an inverted microscope (Olympus IX-83, Olympus, Japan) equipped with a 100X 1.49 NA oil immersion TIRF objective (PlanApi, Olympus). The imaging setup included an s-CMOS camera (ORCA Flash 4.0, Hamamatsu, Japan) and a 488nm laser source. Images were acquired with an exposure time of 200ms, achieving a penetration depth of approximately 70nm. The live cells were activated using 1μg/mL anti-human-CD3 mAb (OKT3) (BD Biosciences, 567107), and time-lapse images of the activated cells were captured.

Image analysis

Image analysis was performed using Fiji Ver1.5. A Gaussian blur operation was performed for cluster analysis on the TIRF images. The blurred image was subtracted from the raw image, thus enhancing the local contrast. The thresholding was performed using the appropriate threshold, resulting in a binary image from which five randomly appearing clusters of 10 pixels (1 pixel= 0.65μm) or larger size were chosen from the cells. The intensity of the chosen clusters was captured over time. The rate of ZAP-70 recruitment to the plasma membrane was determined from the plot of change in EGFP intensity over time. The total cluster number (≥10-pixel size) from the individual cell was determined and analyzed using GraphPad PrismVer9.5.1

**Expression and purification in tandem SH2 domain of ZAP-70**

The tSH2 domains of ZAP-70 and the mutants were overexpressed in *E.coli*-BL21(DE3) cells. The culture was induced with 1 mM IPTG overnight at 18°C. Cells were lysed by sonication in lysis buffer containing 50 mM Tris, pH 8, 200 mM NaCl, 20 mM imidazole, 5 mM β-mercaptoethanol, and 5% glycerol. The tSH2 domain was purified as described previously [1,18]. Briefly, the clear cell lysate was passed through a Ni-NTA affinity column and eluted with imidazole containing elution buffer composed of 50 mM Tris, pH 8, 200 mM NaCl, 500 mM imidazole, 5 mM β-mercaptoethanol, 5% glycerol. The eluate from the Ni-NTA column was further purified using a Q-column followed by gel-filtration chromatography. The purified tSH2 domain was concentrated and stored in 20 mM Tris, pH 8, 150 mM NaCl, 5 mM β-mercaptoethanol, and 5% Glycerol at −80°C.

**Expression and purification in tandem SH2 domain of Syk**

The tSH2 domain of Syk is expressed as GST fusion in *E.coli*-BL21 (DE3) cells as described previously [18]. The cells were lysed by sonication in 1X PBS pH7.4. The cell lysate was passed through the GST affinity column and eluted with L-glutathione containing elution buffer composed of 50 mM Tris, pH 8.2, 10 mM Reduced Glutathione, and 10% Glycerol. The eluted protein was digested overnight using Precision Protease and loaded on a GST column again to remove the cleaved GST tag. The protein was further purified using gel-filtration chromatography. The purified protein was concentrated and stored in 50mM Tris, pH 8, 150mM NaCl, 5mM β-mercaptoethanol, and 10% Glycerol at -80°C.

**Fluorescence Polarization experiments**

The interaction between the tSH2 domain and ITAM-ζ1-$Y_2P$ was quantitatively determined from the steady-state fluorescence anisotropy experiment. The indicated constructs of the tSH2 domain of ZAP-70 were titrated against the 25nM ITAM-ζ1-$Y_2P$ labeled with AlexaFluor® 488. Both the labeled peptide and the tSH2 domain were dissolved in 20mM Tris, pH 8, 150mM NaCl, 5mM β-mercaptoethanol, 5% Glycerol. The fluorescence polarization was recorded using a Hitachi (model F-4500) fluorometer equipped with a polarizer. To test the effect of increasing potassium concentration on the ITAM-ζ1-$Y_2P$ and tSH2 binding, the titration was repeated with various ratios of NaCl and KCl-containing buffers, keeping the total salt concentration at 150mM. The dissociation constant was determined by fitting the curve to the following equation implemented in GraphPad PRISM: $Y = Y_0 + \left[\frac{B_{max} \times X}{K_d + X}\right]$. Where $Y$ is the anisotropy measured in the presence of protein, $Y_0$ is the anisotropy measured for the free ITAM-ζ1-$Y_2P$ peptide, X is the concentration of the protein, $B_{max}$ is the maximum value of anisotropy measured, and $K_d$ is the dissociation constant.

**Thermal Unfolding Assay using Circular Dichroism**

The melting temperature ($T_m$) of the *apo* or *holo* tSH2 domain of ZAP-70 in various KCl concentrations was determined from the thermal unfolding of the proteins measured using a circular dichroism (CD) spectrophotometer (Jasco-J810 spectrophotometer). For the CD experiment, the protein and the peptide were dissolved in 20 mM phosphate buffer pH 7.4 and the various concentrations of NaCl or KCl, keeping the total salt concentration at 150 mM. Two CD spectra of the tSH2 domain were recorded for each experimental condition, one in the *apo* state and one in the *holo* state (1:1 complex of tSH2: ITAM-ζ1-$Y_2P$). For each data set, the spectrum was scanned between 300 to 200 nm at a temperature ranging from 20 to 60°C with an increment of 4°C. The ellipticity data were converted into molar ellipticity using the following equation [19]: $[\theta] = \frac{m_0 \times M}{(10 \times L \times C)}$.

The change in Gibbs free energy of unfolding ($\Delta G_{unfolding}$) for the *apo* or *holo* tSH2 domain was derived assuming the simplest two-step unfolding model [19,20]. The fraction unfolded [U] protein at a given temperature was derived using: $[U] = \frac{(\theta_T - \theta_F)}{(\theta_U - \theta_F)}$. Where $\theta_F$ and $\theta_U$ are the molar ellipticity of the fully folded and unfolded state at 222 nm, $\theta_T$ is the molar ellipticity at a given temperature. The equilibrium constant, $K_{unfolding}$, was calculated using the following equation: $K_{unfolding} = \frac{[U]}{(1-[U])}$ . The $\Delta G_{unfolding}$ was derived from: $\Delta G_{unfolding} = -RTln(K_{unfolding})$, where R= $1.98\times10^{-3}$ kcal $mol^{-1}$ and T= 317K (44°C). Related to Figure 5G.

**Isothermal titration calorimetry**

Malvern PEAQ-ITC was used to determine the affinity of the N-SH2 phosphate binding pocket for the ITAM-ζ1-Y2P peptide at various KCl concentrations. 20μM of ZAP-70 $^{R190A}$tSH2 domain was titrated with increasing ITAM-ζ1-$Y_2$P concentrations in glycerol-free buffer composed of 20 mM HEPES, pH 8.2, 5 mM β-mercaptoethanol and indicated proportion of NaCl and KCl. A stock concentration of 300 μM of ITAM- ζ1-Y2P was loaded in the syringe. Each experiment consisted of nineteen injections of 2 μl each and a delay of 180 s between the injections. The protein solution was stirred at 300 rpm during the titration at 20 ºC. The $K_d$, ΔH, and ΔS were obtained by fitting observed heat exchanged from the titration to a one-site ligand binding model implemented in the MicroCal PEAQ-ITC Analysis Software v1.41.

**Steady-state fluorescence experiments**

Interaction between the tSH2 domain and ITAM-ζ1-$Y_2$P peptide in the steady-state was determined from the change in intrinsic tryptophan fluorescence. The tryptophan fluorescence was measured at $\lambda_{ex}$ 295 nm, and the emission spectrum ($\lambda_{em}$) was scanned between 300 nm to 400 nm at 25 °C. The fluorescence spectra were recorded using a Horiba Duetta spectrophotometer. For each titration, 1 μM tSH2 domain was titrated with various concentrations of ITAM-ζ1-$Y_2$P peptide. The peptide and the protein were dissolved in 20 mM Tris pH 8, 5% glycerol, 5mM β-mercaptoethanol, and different concentrations of NaCl or KCl, keeping the total salt concentration at 150 mM. The normalized fluorescence intensity ($F_0$/F) at 340nm ($\lambda_{em}$) was plotted against the ligand-to-protein molar ratio. Where $F_0$ and F are the intrinsic tryptophan fluorescence in the absence and presence of ligands, respectively.

**Acrylamide Quenching Experiments**

The structure of the tSH2 domain in the open (*apo*) or closed (*holo*) conformation was evaluated from the quenching of intrinsic tryptophan fluorescence by acrylamide at the indicated salt concentration. The tryptophan fluorescence for the *apo* or *holo* tSH2 domain was measured by titrating an increasing acrylamide concentration. The fluorescence emission was scanned between 300-400nm. The fluorescence intensity at 340nm was plotted against the respective acrylamide concentration. The Stern-Volmer quenching constant ($K_{sv}$) was determined from the linear fitting of the data: $\frac{F_0}{F} = 1 + (K_{sv} \times X)$ [1,21]. Where $F_0$ and F are the fluorescence in the absence and presence of acrylamide, respectively. X is the concentration of the acrylamide.

**Pre-steady state kinetics of ITAM-ζ1-Y2P and ZAP-70 tSH2 interaction using Stopped-flow fluorescence spectroscopy**

The binding kinetics of the ZAP-70 tSH2 domain and ITAM-ζ1-$Y_2$P peptides were measured from the change in intrinsic tryptophan fluorescence of the tSH2 domain at 10 °C. The tryptophan fluorescence was recorded with an SFM2000 BioLogic spectrophotometer fitted with a stopped-flow system. The protein and the peptide were dissolved in a buffer containing 20 mM Tris (pH 8.0), 5% glycerol, 5mM β-mercaptoethanol, and different concentrations of NaCl or KCl, as indicated in Figures 6D and 6F. Each kinetic experiment was carried out by mixing 100 nM of tSH2 domain with 15 μM of ITAM-ζ1-$Y_2$P and recorded over 200s (for slow kinetics) or one second (for fast kinetics), respectively. A blank dataset was recorded for each sample by measuring the change in intrinsic tryptophan fluorescence of tSH2 domains upon mixing with buffer. Each data set was normalized against the maximum intensity observed for the sample at t=0. The normalized data set was further corrected by blank subtraction. The observed rate constant ($k_{obs}^{fast}$ or $k_{obs}^{slow}$ ) was derived by fitting to an association kinetics equation using PRISM: $Y = Y_0 + (Y_{max} - Y_0)(1 - e^{-k_{obs}t})$. where $Y_0$ is the intensity at t=0, $Y_{max}$ is the maximum intensity, $k_{obs}$ is the rate constant.

**Table S1**: List of antibodies used

| Antibody | Details | Source | Catalog Number | Lot Number | Dilution |
|---|---|---|---|---|---|
| Anti- Human CD3 Clone: OKT3 | Purified Mouse Anti-Human | BD Pharmagen | 567107 | 3059362 | 1:1000 (Stimulation) |
| Human CD28 Clone:37407 | Mouse Monoclonal | R&D Systems | MAB342 | AEG1124021 | 1:200 (Co-stimulation) |
| Anti-Human IgM | Goat Polyclonal | Sigma-Aldrich | 12386 | 118M4782V | 1:100 (Stimulation) |
| Human LCK Clone:693010 | Mouse Monoclonal | R&D Systems | MAB37041 | CEWU012407B | 1:1000 (Immunoblotting) |
| Anti-Human phospho LCK (Y394) Clone:755103 | Mouse Monoclonal | R&D Systems | MAB7500 | CGED0223041 | 1:1000 (Immunoblotting) |
| Anti-CD3ζ | Rabbit Monoclonal | Cell Signaling Technology | 88083 | 1 | 1:1000 (Immunoblotting) |
| Anti-phospho CD3ζ (Y142) | Rabbit Monoclonal | Cell Signaling Technology | 67748 | 1 | 1:1000 (Immunoblotting) |
| Anti-ZAP-70 (99F2) | Rabbit Monoclonal | Cell Signaling Technology | 2705S | 12 | 1:1000 (Immunoblotting) |
| Anti-phospho ZAP-70 (Y493) | Rabbit Monoclonal | Cell Signaling Technology | 2704S | 10 | 1:1000 (Immunoblotting) |
| Anti-Syk (D3Z1E) | Rabbit Monoclonal | Cell Signaling Technology | 13198S | 9 | 1:1000 (Immunoblotting) |
| Anti-LAT | Rabbit Polyclonal | MyBioSource | MBS2091781 | A202003 | 1:1000 (Immunoblotting) |
| Anti-phospho LAT (Y191) | Rabbit Monoclonal | MyBioSource | MBS82203 | CP1D10A | 1:1000 (Immunoblotting) |
| Anti-GAPDH | Rabbit Monoclonal | BioBharati Life Sciences | BB-AB0060 | 011501 | 1:1000 (Immunoblotting) |
| Anti-PLCγ | Rabbit Monoclonal | Cell Signaling Technology | 5690S | 7 | 1:1000 (Immunoblotting) |

| Antibody | Details | Source | Catalog Number | Lot Number | Dilution |
|---|---|---|---|---|---|
| Anti-phospho PLCγ (Y783) | Rabbit Monoclonal | Cell Signaling Technology | 14008S | 4 | 1:1000 (Immunoblotting) |
| Anti-phospho p44/42 MAPK (T202/Y204) (D13.14.4E) | Rabbit Monoclonal | Cell Signaling Technology | 4370S | 28 | 1:200 (Flow cytometry) |
| Anti-phospho AKT (S473) (193812) | Rabbit Monoclonal | Cell Signaling Technology | 4058T | 30 | 1:200 (Flow cytometry) |
| Anti-rabbit IgG Secondary (HRP) | Goat anti-rabbit | Abcam | Ab6717 | GR267728-27 | 1:2500 (Immunoblotting) |
| Anti-mouse IgG Secondary (HRP) | Goat anti-mouse | Cell Signalling Technology | 7076S | 33 | 1:2500 (Immunoblotting) |
| Anti- total Phosphotyrosine | Rabbit monoclonal | Abcam | AB179530 | 1000218-13 | 1:1000 (Immunoblotting) |
| Anti-rabbit IgG (H+L) Alexa Fluor 488 | Goat anti-Rabbit | Invitrogen | A11008 | 2897813 | 1:100 (Flow cytometry) |
| Anti-rabbit IgG (H+L) Alexa Fluor 647 | Goat anti-Rabbit | Invitrogen | A31573 | 2752586 | 1:100 (Flow cytometry) |
| Anti-Mouse CD3 FITC | Rat Anti-Mouse | BD Pharmagen | 561798 | 1286349 | 1:1000 (Flow cytometry) |
| Anti-Mouse CD4 APC-H7 (Clone: GK1.5) | Rat Anti-Mouse | BD Pharmagen | 560246 | 3352224 | 1:1000 (Flow cytometry) |
| Anti-Mouse CD8a PE | Rat Anti-Mouse | BD Pharmagen | 567630 | 1141239 | 1:1000 (Flow cytometry) |
| Anti-Mouse CD44 PE-Cy7 (Clone: IM7) | Rat Anti-Mouse | BD Pharmagen | 560569 | 1286145 | 1:1000 (Flow cytometry) |
| Anti-Mouse CD25 PE (Clone: 7D4) | Rat Anti-Mouse | BD Pharmagen | 558624 | 1308946 | 1:1000 (Flow cytometry) |
| Anti-Mouse FoxP3 R718 (Clone: MF23) | Rat Anti-Mouse | BD Pharmagen | 567095 | 1253348 | 1:1000 (Flow cytometry) |
| Anti-Mouse CD19 APC-Cy7 | Rat Anti-Mouse | BD Pharmagen | 560143 | 7355677 | 1:1000 (Flow cytometry) |

**Table S2:** Amplitude of calcium signaling measured at t = 540 sec.

| [KCl]mM | [NaCl]mM | Amplitude*(AU) |
|---|---|---|
| 5 | 145 | 2.75±0.28 |
| 10 | 140 | 2.2±0.02 |
| 20 | 130 | 1.48±0.08 |
| 50 | 100 | 1.47±0.187 |
| Clofazimine | | |
| 5 | 145 | 1.13±0.005 |

* The amplitude values represent the mean of three separate experiments and the associated standard deviation

**Table S3:** Rate of decay of PBFI intensity and the intensity of PBFI measured at t = 1200 sec under various salt concentrations and clofazimine treatment.

| [KCl]mM | [NaCl]mM | Rate of Decay (AU/s)* | Relative PBFI ratio * |
|---|---|---|---|
| 5 | 145 | 62.3±3.78 X $10^{-5}$ | 0.56±0.07 |
| 10 | 140 | 44.7±8.26 X $10^{-5}$ | 0.68±0.03 |
| 20 | 130 | 24.3±8.8 X $10^{-5}$ | 0.73±0.11 |
| 50 | 100 | 34.7±3.68 X $10^{-5}$ | 0.77±0.03 |
| Clofazimine | | | |
| 5 | 145 | 1.53±0.55 X $10^{-5}$ | 0.98±0.008 |

* The values represent the mean of three separate experiments measured at 1200 sec and the associated standard deviation.

**Table S4:** Binding parameters of ITAM-ζ1-$Y_2$P and the C-SH2 phosphate binding pocket determined from fluorescence polarization experiment.

| [KCl]mM | [NaCl]mM | $K_{d1}$ (nM)* | $\Delta G^1_{Binding}$ (kcal/mol)* | $\Delta\Delta G^1_{Binding}$ (kcal/mol)* |
|---|---|---|---|---|
| 0 | 150 | 4.52±0.93 | -11.16±0.12 | -- |
| 5 | 145 | 4.44±0.69 | -11.17±0.1 | -0.006±0.071 |
| 20 | 130 | 4.34±0.42 | -11.17±0.06 | -0.015±0.074 |
| 50 | 100 | 4.53±0.66 | -11.15±0.087 | 0.006±0.094 |

* The values represent the mean of three separate experiments and the associated standard deviation

**Table S5:** Binding parameters of ITAM-ζ1-$Y_2$P and the N-SH2 phosphate binding pocket determined from isothermal titration calorimetry experiment.

| [KCl ]mM | [NaCl] mM | $K_{d2}$ (µM)* | $\Delta G^2_{Binding}$ (kcal/mol)* | TΔS (kcal/mol)* | ΔH (kcal/mol)* | N | $\Delta\Delta G^2_{Binding}$ (kcal/mol)* |
|---|---|---|---|---|---|---|---|
| 0 | 150 | 6.33±0.34 | -6.95±0.031 | 5.6±0.1 | -11.5±0.17 | 0.9 | -- |
| 5 | 145 | 9.77±0.61 | -6.69±0.037 | 4.75±0.12 | -10.4±0.21 | 0.9 | 0.25±0.012 |
| 20 | 130 | 20.93±0.82 | -6.25±0.023 | 3.04±0.08 | -9.35±0.196 | 0.9 | 0.69±0.009 |
| 50 | 100 | 26.17±0.85 | -6.12±0.02 | 2.64±0.14 | -8.77±0.11 | 0.9 | 0.82±0.027 |

* The values represent the mean of three separate experiments and the associated standard deviation

**Table S6:** Change in Gibbs free energy for unfolding ($\Delta G_{unfolding}$) measured from the thermal denaturation of *apo* and *holo* tSH2 domain of ZAP-70 using CD spectroscopy**.**

| *Apo* | | | |
|---|---|---|---|
| [KCl] mM | [NaCl] mM | Melting Temperature $T_m$(°C) | $\Delta G_{unfolding}$ (kcal/mol) |
| 0 | 150 | 39.13±1.375 | -0.405±0.135 |
| 1 | 149 | 40.02±0.57 | -0.431±0.105 |
| 3 | 147 | 40.24±1.065 | -0.862±0.175 |
| 7 | 143 | 40.58±0.565 | -0.550±0.068 |
| 10 | 140 | 38.04±0.53 | -0.528±0.237 |
| 50 | 100 | 37.54±1.51 | -0.407±0.129 |
| 100 | 50 | 37.29±0.585 | -0.963±0.099 |
| 150 | 0 | 38.27±0.645 | -0.781±0.050 |
| *Holo* | | | |
| [KCl] mM | [NaCl] mM | Melting Temperature $T_m$(°C) | $\Delta G_{unfolding}$ (kcal/mol) |
| 0 | 150 | 47.79±0.27 | 1.083±0.033 |
| 1 | 149 | 46.88±0.295 | 0.880±0.012 |
| 3 | 147 | 46.45±0.3 | 0.770±0.071 |
| 7 | 143 | 45.69±0.375 | 0.736±0.005 |
| 10 | 140 | 45.67±0.63 | 0.651±0.021 |
| 50 | 100 | 45.61±0.2 | 0.570±0.035 |
| 100 | 50 | 45.33±0.195 | 0.493±0.064 |
| 150 | 0 | 44.37±0.28 | 0.434±0.087 |

* $\Delta G_{unfolding}$ is calculated at T= 317K (44°C).

**Table S7:** Stern-Volmer quenching constant ($K_{sv}$) for the *apo* and *holo* tSH2 domain of ZAP-70 at indicated salt concentration.

| Apo | | |
|---|---|---|
| **[KCl]mM** | **[NaCl]mM** | **$K_{sv}$ (µM$^{-1}$)*** |
| 0 | 150 | 0.052±0.004 |
| 5 | 145 | 0.062±0.004 |
| 20 | 130 | 0.066±0.004 |
| 50 | 100 | 0.068±0.001 |
| *Holo* | | |
| **[KCl]mM** | **[NaCl]mM** | **$K_{sv}$ (µM$^{-1}$)*** |
| 0 | 150 | 0.0164±0.001 |
| 5 | 145 | 0.0278±0.001 |
| 20 | 130 | 0.03±0.001 |
| 50 | 100 | 0.042±0.002 |

* The values represent the mean of three separate experiments and the associated standard deviation

**Table S8:** Observed rate ($k_{obs}$) for the ZAP-70 tSH2 domain and ITAM-ζ1-$Y_2$P binding measured at indicated salt concentration.

| Fast Binding ($k_{obs}^{Fast}$) | | |
|---|---|---|
| **[KCl]mM** | **[NaCl]mM** | **$k_{obs}$ ($s^{-1}$)*** |
| 0 | 150 | 21.74±2.52 |
| 5 | 145 | 25.68±2.79 |
| 20 | 130 | 25.07±4.61 |
| Slow Binding ($k_{obs}^{Slow}$) | | |
| 0 | 150 | 0.23±0.007 |
| 5 | 145 | 0.132±0.013 |
| 20 | 130 | 0.1±0.0003 |

* The values represent the mean of three separate experiments and the associated standard deviation

**Table S9:** Dissociation constant ($K_d$) determined from the titration of ITAM-ζ1-$Y_2$P and the tSH2 domain of Syk by measuring the changes in intrinsic tryptophan fluorescence.

| [KCl]mM | [NaCl]mM | $K_d$ (nM)* |
|---|---|---|
| 0 | 150 | 97.54±18.5 |
| 5 | 145 | 86±10.4 |
| 20 | 130 | 84.3±12.1 |
| 50 | 100 | 82.67±12.01 |

* The values represent the mean of three separate experiments and the associated standard deviation

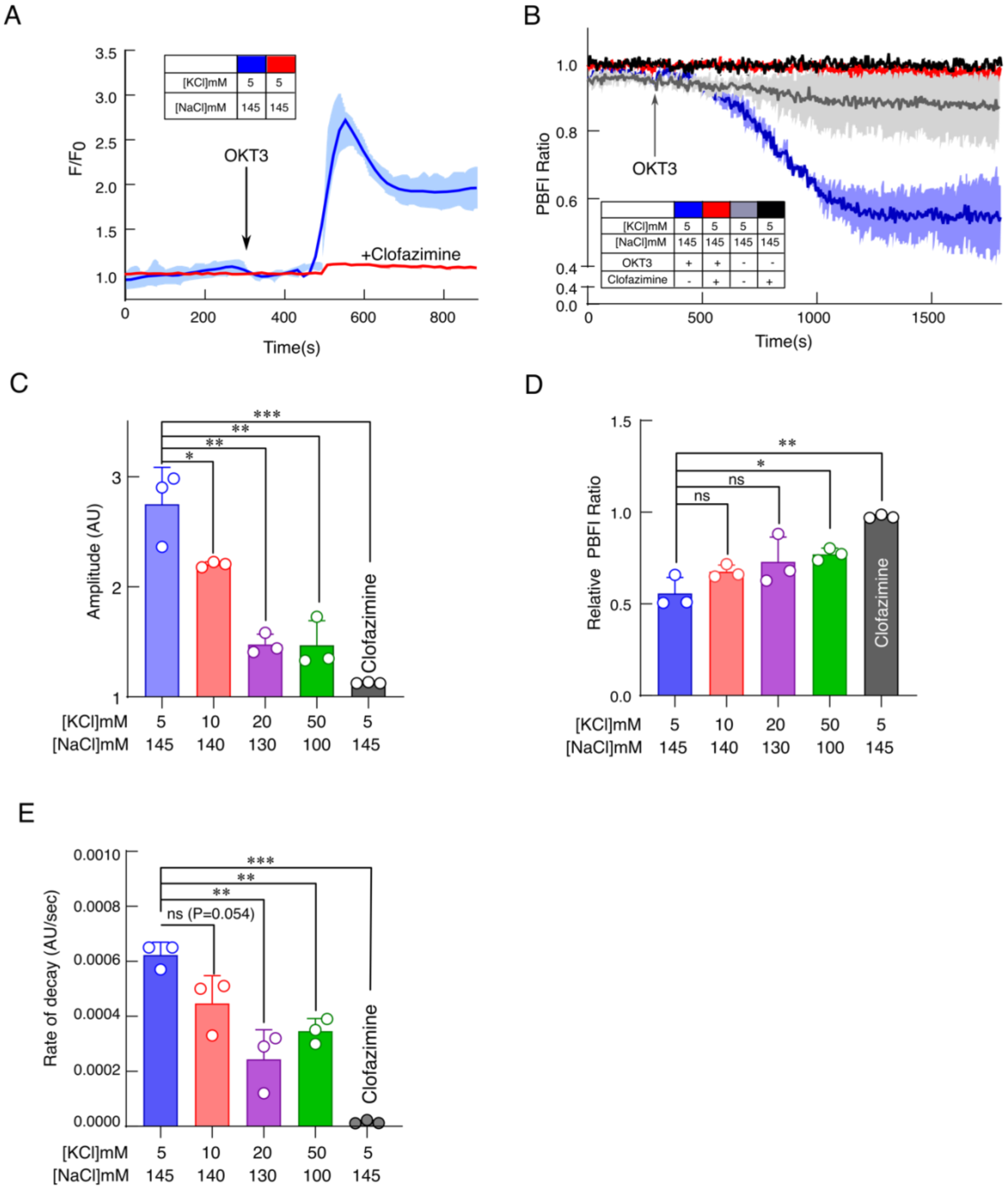
A
F/F0
3.5
3.0
2.5
2.0
1.5
1.0
0
200
400
600
800
Time(s)
OKT3
+Clofazimine
[KCl]mM 5 5
[NaCl]mM 145 145
B
PBFI Ratio
1.0
0.8
0.6
0.4
0.0
0
500
1000
1500
Time(s)
OKT3
[KCl]mM 5 5 5 5
[NaCl]mM 145 145 145 145
OKT3 + + - -
Clofazimine - + - +
C
Amplitude (AU)
3
2
1
*
**
**
***
Clofazimine
[KCl]mM 5 10 20 50 5
[NaCl]mM 145 140 130 100 145
D
Relative PBFI Ratio
1.5
1.0
0.5
0.0
ns
ns
*
**
Clofazimine
[KCl]mM 5 10 20 50 5
[NaCl]mM 145 140 130 100 145
E
Rate of decay (AU/sec)
0.0010
0.0008
0.0006
0.0004
0.0002
0.0000
ns (P=0.054)
**
**
***
Clofazimine
[KCl]mM 5 10 20 50 5
[NaCl]mM 145 140 130 100 145

**Figure S1: Potassium channel inhibitor, clofazimine, prevents TCR-dependent potassium efflux and attenuates calcium signaling.**

A) Calcium flux was measured from the plot of normalized Fluo4 fluorescence intensity as a function of time (seconds) on stimulation of Jurkat E6.1 T-cells in the absence (blue solid line) and presence of clofazimine (red solid line). The T-cells were stimulated with anti-CD3 antibody (OKT3), at the indicated time. The fluorescence intensity for the activated (F) T-cells is normalized against the inactive ($F_o$) cells. The solid lines represent the mean from three independent experiments, and the area fill denotes the standard deviation.

B) The potassium efflux is measured from the plot of PBFI fluorescence intensity as a function of time in the Jurkat E6.1 T-cells in the presence (red) and absence (blue) of clofazimine. The T-cells were activated with an anti-CD3 antibody (OKT3), at the indicated time. The fluorescence intensity from uninduced cells is represented as solid black and gray lines. The solid lines represent the mean from three independent experiments, and the area fill denotes the standard deviation.

C) The bar graph represents the amplitude of calcium signaling in activated Jurkat E6.1 T-cells under indicated experimental conditions. From left to right $P = 0.0488$, $P = 0.0032$, $P = 0.0054$, $P = 0.0011$.

D) Bar graph representing the rate of potassium efflux in activated Jurkat E6.1 T-cells under indicated experimental conditions. From left to right $P = 0.0547$, $P = 0.0052$, $P = 0.0017$, $P < 0.0001$.

E) Bar graph representing the mean intensity ratio of PBFI recorded in activated Jurkat E6.1 T-cells at 1200 seconds, as shown in Figures 1C and S1B. From left to right $P = 0.0944$, $P = 0.1356$, $P = 0.0162$, $P = 0.0012$)

Statistical analysis of two-tailed Student's t test performed. In the bar plots (panel C-D), the data represent mean $\pm$ SD (ns= not significant; *$P<0.05$; **$P<0.01$; ***$P<0.001$; ****$P<0.0001$). All data were plotted using GraphPad PrismVer9.5.1.

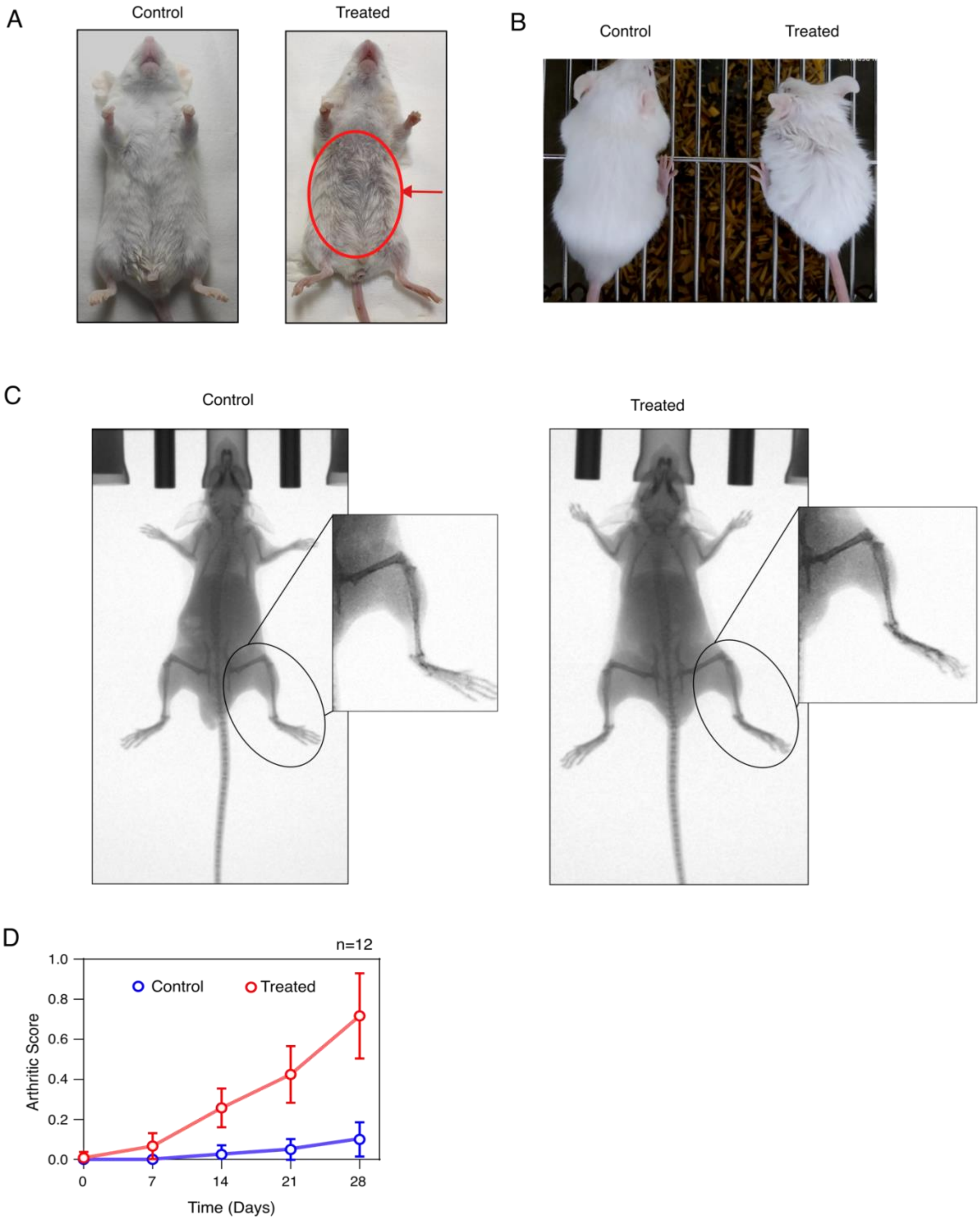
A
Control
Treated
B
Control
Treated
C
Control
Treated
D
n=12
Control
Treated
Arthritic Score
1.0
0.8
0.6
0.4
0.2
0.0
0
7
14
21
28
Time (Days)

**Figure S2: Elevated serum potassium induces higher arthritis-like scores in juvenile mice**

A) and B) Representative images showing the loss of body hair in the treated mice (on the right) compared to the untreated control (on the left) mice.

C) Representative full-body X-ray images of control (on the left) and treated (on the right) mice. The zoom-in section of the knee joint region is shown in the inset. In the treated mice, the following observations are made: joint space narrowing, bone erosions, increased opacity, and soft tissue swelling. The X-ray images indicate progressive joint destruction in the treated mice.

D) Arthritic score is plotted as a function of time (in days) observed in control (blue) and treated (red) groups of mice (n=12). The solid line is the guiding line. Each data point represents the mean ± SD.

All data were plotted using GraphPad PrismVer9.5.1. The schematics and icons were made using Inkscape Ver1.4.

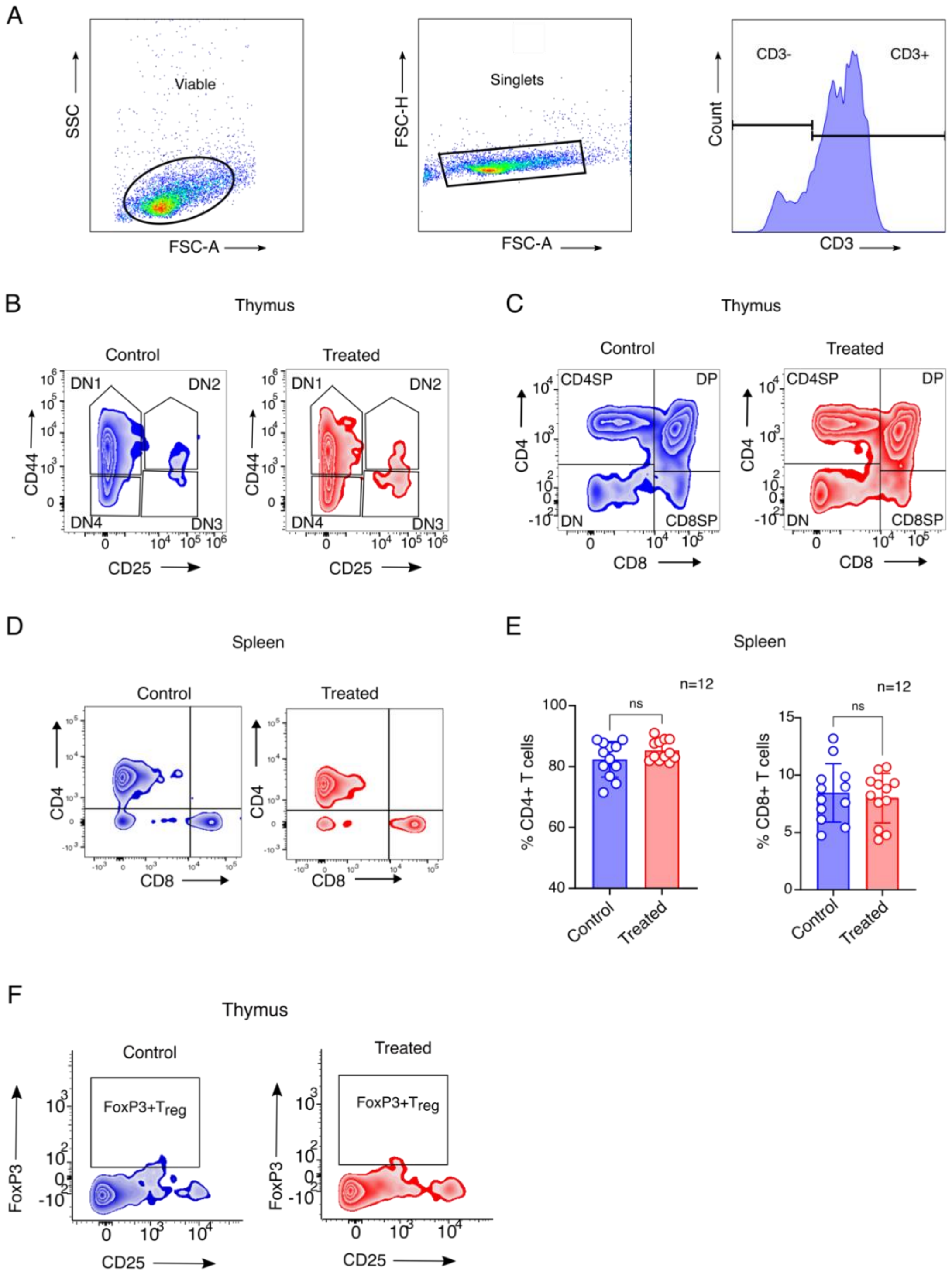

A
Viable
SSC
FSC-A
Singlets
FSC-H
FSC-A
CD3-
CD3+
Count
CD3
B
Thymus
Control
Treated
DN1
DN2
DN4
DN3
CD44
CD25
C
Thymus
Control
Treated
CD4SP
DP
DN
CD8SP
CD4
CD8
D
Spleen
Control
Treated
CD4
CD8
E
Spleen
n=12
ns
% CD4+ T cells
Control
Treated
n=12
ns
% CD8+ T cells
Control
Treated
F
Thymus
Control
Treated
FoxP3+Treg
FoxP3
CD25

**Figure S3: Immune profiling of control and hyperkalemic juvenile mice**

A) A representative gating strategy used for flow cytometric analysis of experimental animal tissues. The SSC and FSC-A/H indicate side and forward scatter (area/height), respectively.

B) Representative contour plot from the flow cytometric analysis of thymic $CD3^{+}$ double negative (DN) T cells in control (blue) and treated (red) groups of mice. The cells were stained with anti-CD3 (FITC), anti-CD25 (PE), and anti-CD44 (PE-Cy7) mAb to study the different Double Negative (DN) stages of the developing T thymocytes.

C) and D) Representative contour plot from the flow cytometric analysis of $CD3^{+}$ T cells in the thymus and spleen, respectively, for the control (blue) and treated (red) groups of mice. The cells were stained with anti-CD3 (FITC), anti-CD4 (APC-H7), and anti-CD8a (PE) mAb to study the Double Negative, Double Positive, and Single Positive stages in the thymus (B) and the Single Positive stages in the spleen (C).

E) The bar graph compares the percentage of splenic total $CD4^{+}$ (on the left) and $CD8^{+}$ (on the right) T-cells in the control (blue) and treated (red) groups of mice (n=12). The data represents mean $\pm$ SD. $CD4^{+}$ T-cell $P=0.1436$; $CD8^{+}$ T-cells $P=0.6394$; Statistical analysis of two-tailed Student's t-test performed. (ns= not significant; $^{*}P<0.05$; $^{**}P<0.01$; $^{***}P<0.001$; $^{****}P<0.0001$).

F) Representative flow cytometric analysis to determine the frequencies of $CD4^{+}$regulatory T-cells ($T_{reg}$) in the thymus. The $T_{reg}$-cells are labeled with anti-CD3 (FITC), anti-CD4 (APC-H7), anti-CD25 (PE), and anti-FoxP3 (R718) mAb.

All data were plotted using GraphPad PrismVer9.5.1. The flow cytometry data were analyzed using FlowJo Ver8.

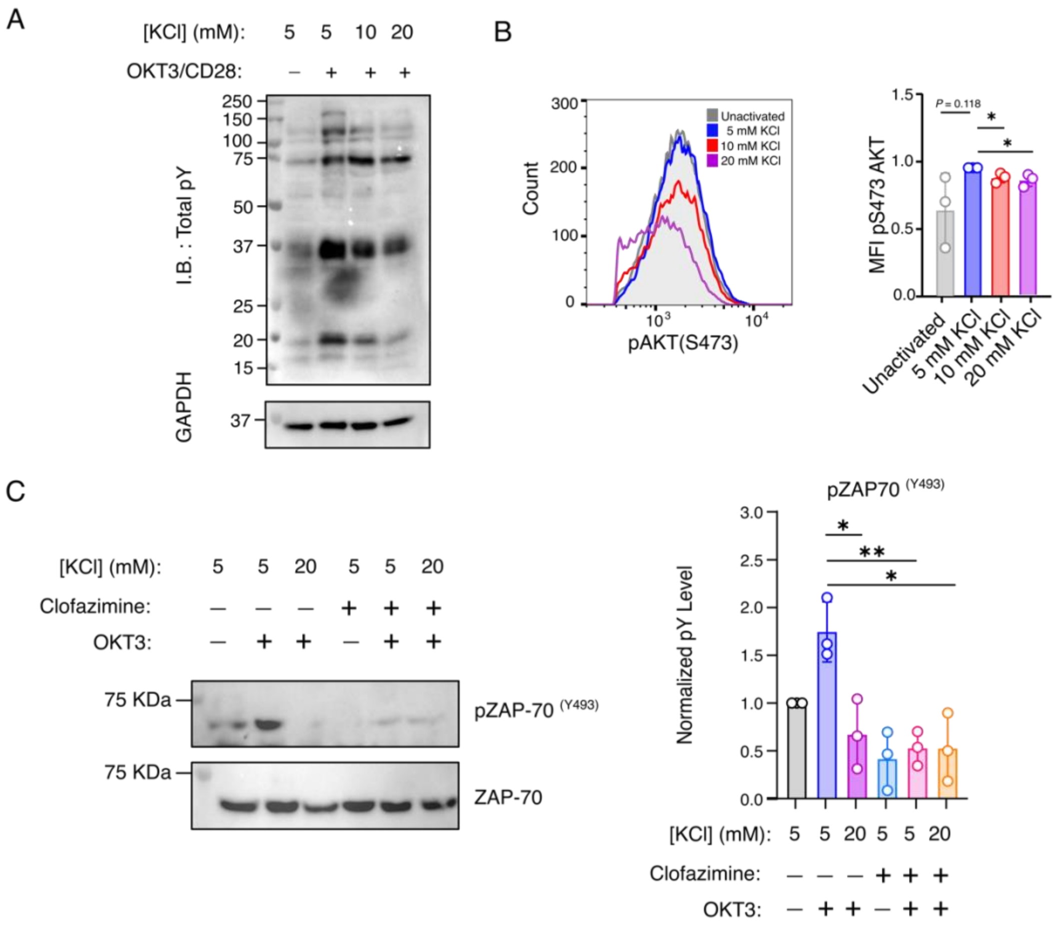


**Figure S4: Clofazimine treatment attenuates ZAP-70 activation following TCR stimulation.**

A) Representative immunoblot analysis of total phosphotyrosine levels in Jurkat E6.1 T-cells in the indicated experimental conditions. Anti-GAPDH mAb staining is the loading control.

B) Representative flow cytometry histograms (on the left) of phospho-AKT (S473) in stimulated Jurkat E6.1 T-cells at the indicated extracellular potassium concentration. Adjacent bar graphs (on the right) show the fold change in AKT phosphorylation at the indicated extracellular potassium concentration (n=3). From left to right, *P*=0.118; *P*= 0.0268; *P*= 0.0286.

C) Representative immunoblot analysis of ZAP-70 activation in Jurkat E6.1 T-cells at the indicated experimental conditions. The cells were activated with OKT3/anti-CD28 for 5mins at $37^0$C. The ZAP-70 activation was determined using ZAP-70-specific anti-pY493 mAb and anti-ZAP70 mAb used as a loading control. The bar graph (on the right) shows the relative phosphorylation level of ZAP-70 Y493 at the indicated experimental conditions (n=3). From left to right, *P*=0.0182; *P*=0.0043; *P*=0.0115.

Statistical analysis of two-tailed Student's t-test performed. Central values and error bars in the bar plot represents mean ± SD (ns= not significant; **P*<0.05; ***P*<0.01; ****P*<0.001; *****P*<0.0001). All data were plotted using GraphPad PrismVer9.5.1. The flow cytometry data were analyzed using FlowJo Ver8. The schematics and icons were made using Inkscape Ver1.4.

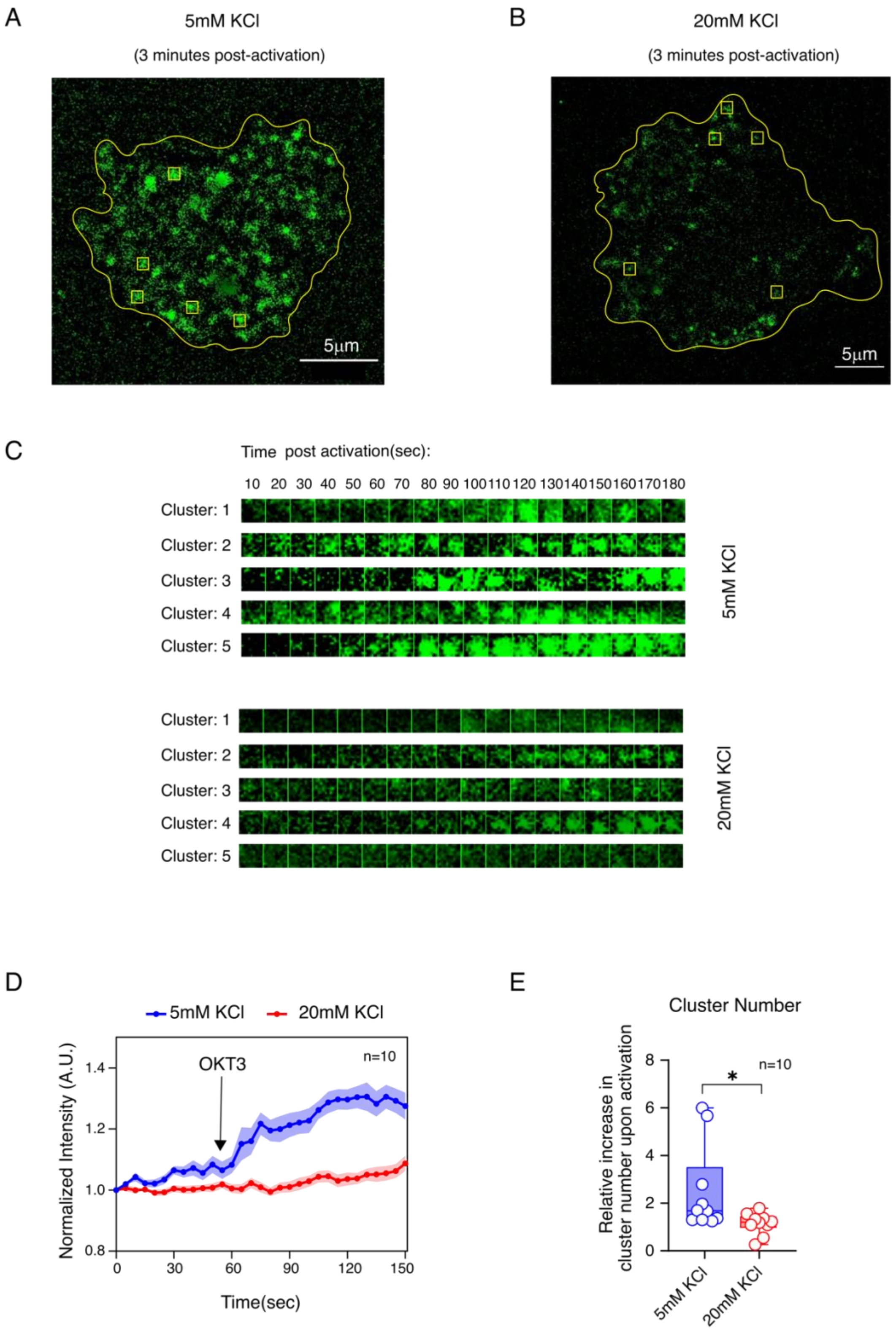

A
5mM KCl
(3 minutes post-activation)
5μm
B
20mM KCl
(3 minutes post-activation)
5μm
C
Time post activation(sec):
10 20 30 40 50 60 70 80 90 100 110 120 130 140 150 160 170 180
Cluster: 1
Cluster: 2
Cluster: 3
Cluster: 4
Cluster: 5
5mM KCl
20mM KCl
D
5mM KCl
20mM KCl
OKT3
n=10
Normalized Intensity (A.U.)
Time(sec)
E
Cluster Number
Relative increase in cluster number upon activation
n=10
*
5mM KCl
20mM KCl

**Figure S5: Membrane recruitment of ZAP-70 by TIRF microscopy**

A) and B) Representative live-cell images of Jurkat P116 T-cell showing the recruitment of ZAP-70 at three minutes post-activation with OKT3/anti-CD28 in the presence of 5 mM (A) or 20 mM (B) extracellular KCl. The cell boundary is marked with a yellow line, and the yellow box indicates five random clusters.

C) Time-lapse montage of five ZAP-70 clusters formed after activating with OKT3/CD28 mAb in Jurkat P116 cells stably expressing ZAP-70-EGFP in the presence of 5mM (top panel) and 20mM (bottom panel) extracellular KCl concentration.

D) The plot of mean EGFP intensity as a function of time measured from ten Jurkat P116 cells stably expressing ZAP-70-EGFP. The arrow indicates the time point when OKT3/ anti-CD28 mAb is added. Each point represents the mean of the average intensity of five ZAP-70 clusters (size: 10 pixels, 1pixel= 0.65μm) from each cell (n=10 cells), and the error bar denotes the std error of the mean. The red and blue solid lines are guiding lines.

E) Box plot showing the relative increase in the number of clusters in P116 cells stably expressing ZAP-70-EGFP at 3 minutes post-activation with OKT3/anti-CD28 (n=10 cells). $P$=0.034.

A statistical analysis of two-tailed Students' t-tests was performed. Data in panel E represents mean ± SD (ns= not significant; *$P$<0.05; **$P$<0.01; ***$P$<0.001; ****$P$<0.0001). The data in panel E is plotted using GraphPad PrismVer9.5.1. The image analysis was done using Fiji Ver 1.54m. The schematics and icons were made using Inkscape Ver1.4.

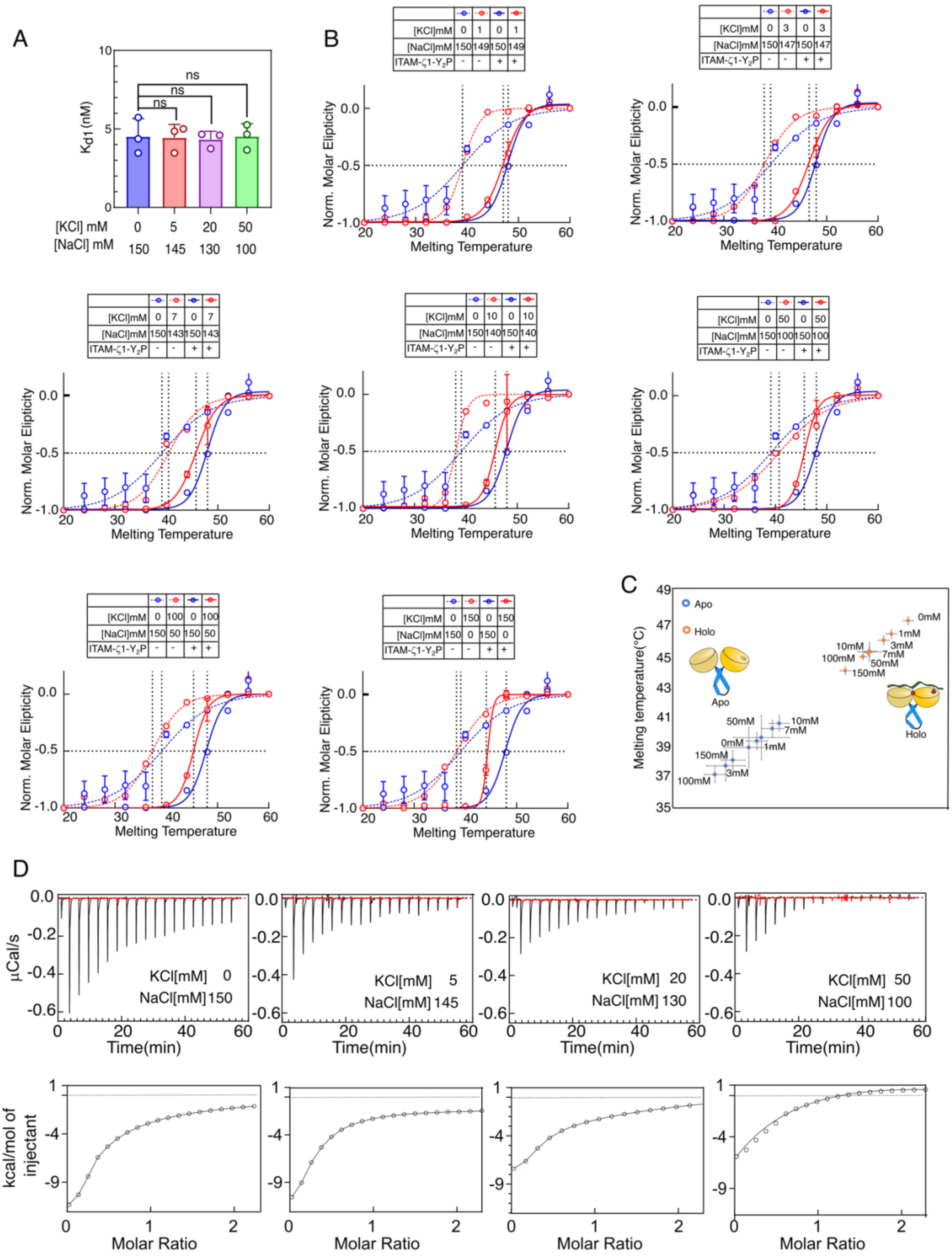
A
ns
ns
ns
Kd1 (nM)
10
5
0
[KCl] mM 0 5 20 50
[NaCl] mM 150 145 130 100
B
[KCl]mM 0 1 0 1
[NaCl]mM 150 149 150 149
ITAM-ζ1-Y2P - - + +
[KCl]mM 0 3 0 3
[NaCl]mM 150 147 150 147
ITAM-ζ1-Y2P - - + +
[KCl]mM 0 7 0 7
[NaCl]mM 150 143 150 143
ITAM-ζ1-Y2P - - + +
[KCl]mM 0 10 0 10
[NaCl]mM 150 140 150 140
ITAM-ζ1-Y2P - - + +
[KCl]mM 0 50 0 50
[NaCl]mM 150 100 150 100
ITAM-ζ1-Y2P - - + +
[KCl]mM 0 100 0 100
[NaCl]mM 150 50 150 50
ITAM-ζ1-Y2P - - + +
[KCl]mM 0 150 0 150
[NaCl]mM 150 0 150 0
ITAM-ζ1-Y2P - - + +
Norm. Molar Elipticity
0.0
-0.5
-1.0
20 30 40 50 60
Melting Temperature
C
Apo
Holo
Melting temperature(°C)
49 47 45 43 41 39 37 35
0mM
1mM
3mM
10mM
7mM
100mM
50mM
150mM
Apo
Holo
50mM
10mM
7mM
0mM
1mM
150mM
3mM
100mM
D
µCal/s
0.0
-0.2
-0.4
-0.6
KCl[mM] 0
NaCl[mM] 150
KCl[mM] 5
NaCl[mM] 145
KCl[mM] 20
NaCl[mM] 130
KCl[mM] 50
NaCl[mM] 100
0 20 40 60
Time(min)
kcal/mol of injectant
1
-4
-9
0 1 2
Molar Ratio

**Figure S6: Potassium prevents ITAM-ζ1-$Y_2P$ and ZAP-70 tSH2 interaction in a concentration dependent manner**

A) The bar graph represents the dissociation constant ($K_{d1}$) for the ITAM-ζ1-$Y_2P$ binding to the C-SH2 phosphate-binding pocket. The $K_{d1}$ *was* obtained from the titration of ITAM-ζ1-$Y_2P$ and $^{R39A}$tSH2 domains by fluorescence polarization experiment. From left to right, $P$=0.5628, $P$=0.9365, $P$=0.1531.

B) Thermal denaturation profiles of *apo* (broken lines) and *holo* (solid line) ZAP-70 tSH2 domain were measured using CD spectroscopy at the indicated potassium concentration. The lines represent the fitting to the Boltzmann sigmoidal equation. The intersect between the black-dotted vertical and horizontal lines indicates the $T_m$.

C) Melting temperature ($T_m$) for the *apo* (blue circle) and *holo* (orange circle) tSH2 domain of ZAP-70 measured at increasing KCl concentration is plotted. The melting temperature is derived from the thermal denaturation profile measured from CD spectroscopy (n=3).

D) ITC titration of ITAM-ζ1-$Y_2P$ and ZAP-70 tSH2 construct bearing R190A mutation. For each titration, 20 µM of tSH2 was titrated with 300 µM of ITAM-ζ1-$Y_2P$. *Top panel*: Black lines represent protein and ligand titration, and the *red line* represents buffer-to-buffer titration. *Bottom panel*: The *solid line* represents the fitting to the one-site binding model.

A statistical analysis of two-tailed Students' t-tests was performed. Each data point represent mean ± SD (ns= not significant; *$P$<0.05; **$P$<0.01; ***$P$<0.001; ****$P$<0.0001). All data were plotted using GraphPad PrismVer9.5.1. Fitting of the ITC data was done using Malvern PEAQ ITC Analysis Software (v 1.41). The schematics and icons were made using Inkscape Ver1.4.

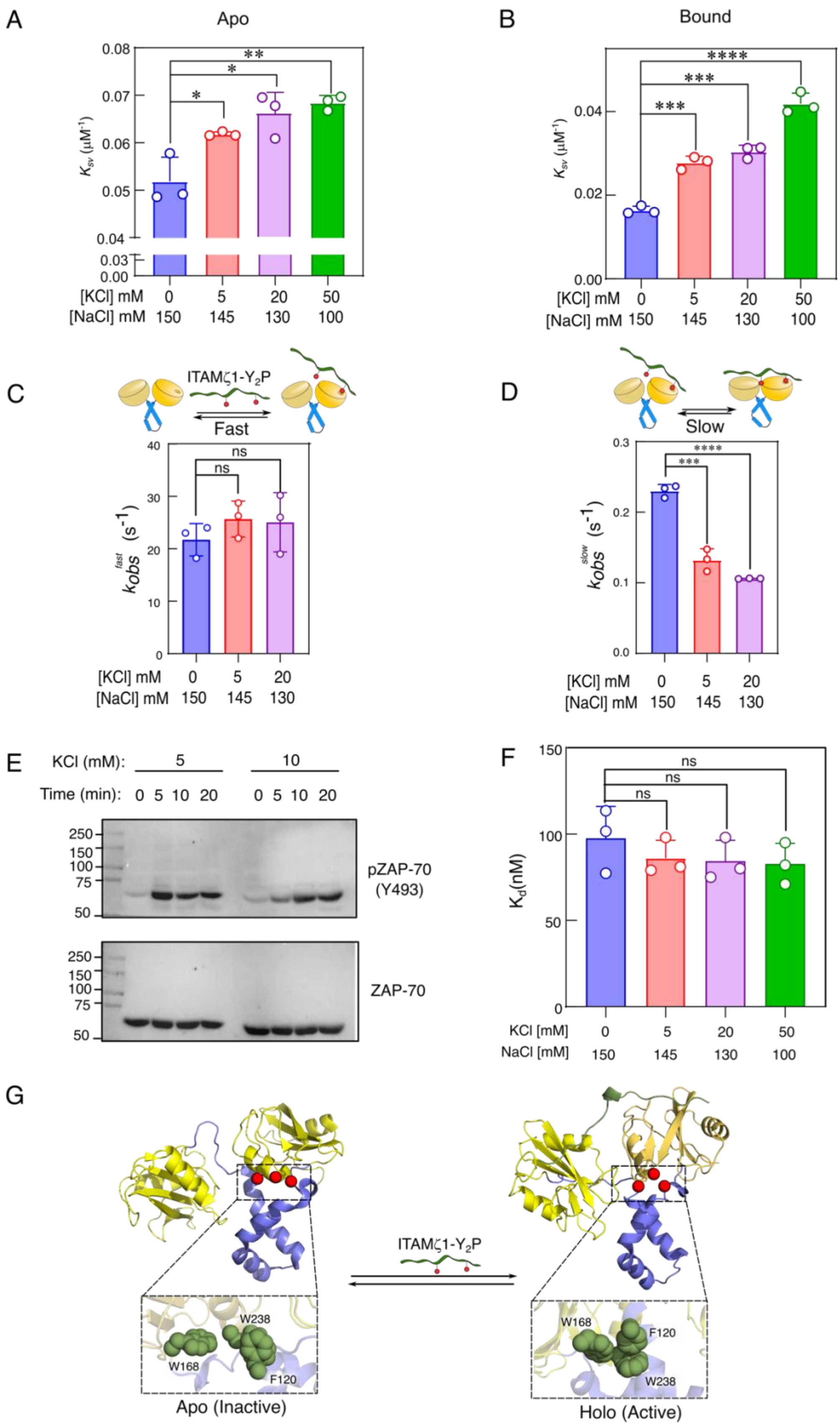

A
Apo
$K_{sv}$ (μM⁻¹)
[KCl] mM 0 5 20 50
[NaCl] mM 150 145 130 100
B
Bound
C
ITAMζ1-Y2P
Fast
$k_{obs}^{fast}$ (s⁻¹)
ns
D
Slow
$k_{obs}^{slow}$ (s⁻¹)
E
KCl (mM): 5 10
Time (min): 0 5 10 20 0 5 10 20
pZAP-70 (Y493)
ZAP-70
F
Kd(nM)
ns
KCl [mM] 0 5 20 50
NaCl [mM] 150 145 130 100
G
W238
W168
F120
Apo (Inactive)
Holo (Active)

**Figure S7: High potassium reduces the rate of structural transition of tSH2 domains from an open to a closed state**

A) and B) The bar graph represents the Stern-Volmer quenching constant ($K_{sv}$) for the *apo* and *holo* tSH2 domain at the indicated salt concentration, respectively (n=3). From left to right *P*=0.0284, *P*=0.0230, *P*=0.0058, *P*=0.0004, *P*=0.0002, *P*=<0.0001.

C) and D) The bar graph represents the rate of change in intrinsic tryptophan fluorescence ($K_{obs}^{fast}$ and $K_{obs}^{slow}$) of the tSH2 domain upon ITAM-ζ1-$Y_2$P binding at the indicated salt concentration. The $K_{obs}^{fast}$ and $K_{obs}^{slow}$ are obtained from the pre-steady state experiments described in Figure 6D and F. (n=3). From left to right *P*=0.2125, *P*=0.4207, *P*<0.0001, *P*=0.0007.

E) A representative immunoblot of ZAP-70 Y493 autophosphorylation upon stimulating Jurkat cells in the presence of indicated extracellular KCl concentration.

F) The bar graph representing the dissociation constant ($K_d$) of the tSH2 domain of Syk and ITAM-ζ1-$Y_2$P determined in the presence of indicated salt composition. The $K_d$ was calculated from the change in tSH2 domain intrinsic tryptophan fluorescence during the titration of ITAM-ζ1-$Y_2$P. From left to right *P*=0.4001, *P*=0.3593, *P*=0.3078.

G) Structure of the tSH2 domain of Syk in the *apo* (PDB ID: 1A81) and *holo* (PDB ID: 4FL2) states. The conformation of the aromatic residues is shown in the inset.

Panels A- D, and F, a statistical analysis of two-tailed Students' t-tests was performed. Each data point represents mean ± SD (ns= not significant; *$P$<0.05; **$P$<0.01; ***$P$<0.001; ****$P$<0.0001). All data were plotted using GraphPad PrismVer9.5.1. The schematics and icons were made using Inkscape Ver1.4.
